# Back to the future 2: the implications of germplasm structure on the balance between short and long-term genetic gain in a changing target population of environments

**DOI:** 10.1101/2025.09.01.673542

**Authors:** Frank Technow, Dean Podlich, Mark Cooper

## Abstract

Plant breeding operates within a highly complex genetic landscape determined by gene effects emerging through biological networks and their interactions with the environment. This environment is not constant but subject to short-term fluctuations and long-term shifts. This significantly complicates the task of plant breeders in finding a balance between adapting their germplasm for the short- and long-term. Here we build on previous work of us that investigated the implications of genetic complexity on breeding program design, by adding an environmental dimension in the form of the *E(NK)* model to the simulation framework. We found that the addition of environmental interactivity and change creates greater uncertainty associated with pursuing any specific selection trajectory, as compared to a static environment. This advantages preserving genetic variability and genetic landscape exploration over quickly exposing additive variation by constraining genetic space around a particular and temporary local optimum. Nonetheless, we found that also in a dynamically changing environment, a structure in which several breeding programs explore genetic space while maintaining constant germplasm exchange, finds the best balance between short and long-term objectives, as opposed to isolated programs or one large undifferentiated program, which exclusively emphasize short respectively long-term objectives. We furthermore highlight the difficulty of exchanging germplasm to restore genetic variability with non-stationary and germplasm context dependent genetic effects. In summary we found that also with addition of environmental complexity and change, the structural features that characterized breeding operations hitherto and allowed them to navigating biological complexity apply. Namely the necessity to constraining genetic space in order for heritable additive variation to emerge. We end by arguing that optimal breeding program design depends on the level of genetic and environmental complexity, which should be appropriately reflected when modeling the long-term behavior of selection programs and the implications of specific interventions into these.

## Introduction

The success and productivity of agricultural crops, like all plant species, depend on the system of biotic and abiotic, natural and man-made, environmental features they are embedded in (Comstock and Moll 1963; Hammer *et al*. 2006). Examples are nutrient and water availability, soil properties and preparation, weed, pest and pathogen pressure as well as the various weather factors like temperature and solar radiation (Clements 1964; Hammer *et al*. 2020; Xing *et al*. 2023). (We use the term “environment” to refer to all extraneous factors influencing plant growth and development, including management practices.) The combinations of the various environmental factors are virtually endless. Individual environments are therefore often grouped into a smaller number of discrete environment types (ET), in which environmental conditions are relatively homogeneous (Gauch Jr. and Zobel 1997; Chenu 2015). The target population of environments (TPE) describes the relative frequencies of the different ET within a given crop growing region (Comstock 1977; Cooper and Podlich 2002). Maximizing crop productivity across a TPE is one of the main concerns of farmers, agronomists and plant breeders alike (Cooper *et al*. 2022). Because of the diversity of a typical TPE, no single plant genotype can be optimally adapted to all ETs found therein, leading to a phenomenon called genotype by environment interaction (GxE), i.e., the differential relative performance of genotypes across environments. GxE is one of the most fundamental problems of plant breeding and considerable efforts have been and are undertaken to characterize, mitigate and predict its effects (e.g., Comstock and Moll 1963; Cooper and DeLacy 1994; Piepho 1998; Elias *et al*. 2016; van Eeuwijk *et al*. 2016; Cooper *et al*. 2022). The TPE itself, however, is continuously changing as a result of processes like climate change (Chapman *et al*. 2012; Peng *et al*. 2020; Cooper *et al*. 2021) and the evolution of weed, pest and pathogens species (McCann 2020), as well as policy, cultural and economic shifts that affect crop management practices, e.g., the availability of mineral and organic nitrogen fertilizer (Kanter *et al*. 2020; Moore *et al*. 2023). Thus, not only is universally optimal adaptation across a TPE virtually impossible, but even what constitutes optimal adaptation is an ever moving target, which is further complicating the task of plant breeders. Ensuring the adaptability of germplasm to a changing TPE, particularly under the pressure of climate change, is therefore one of the main concerns of plant breeders and scientists alike (Kusmec *et al*. 2021; Xiong *et al*. 2022). Different approaches to this challenge have been proposed, such as mining and utilization of gene bank resources (Mayer *et al*. 2020), accelerated variety development (Atlin *et al*. 2017), improved germplasm access and exchange (Galluzzi *et al*. 2020), selection under conditions mimicking future environments in silico and in situ (Cooper and Messina 2023), design of climate resilient ideotypes (Semenov and Stratonovitch 2013) and more recently also the use of gene editing technology (Karavolias *et al*. 2021). However, like natural populations, plant breeding germplasm has an inherent ability to adapt to environmental change through the time tested application of recurrent selection (Becklin *et al*. 2016; Gao *et al*. 2023). One of the most consequential decisions breeders have to make is about their breeding program structure, i.e., whether or not to stratify their germplasm into more or less independent sub-populations or programs and how much if any germplasm exchange between those to allow (Baker and Curnow 1969; Podlich and Cooper 1999; Cooper *et al*. 2014; Technow *et al*. 2021; Yabe *et al*. 2016; Ramasubramanian and Beavis 2021). We use the collective term germplasm to refer to the standing genetic variation for traits within the reference population of genotypes that is utilized in combination with the chosen breeding program structure to create and evaluate new genotypes to achieve the breeding program objectives. In this study we aim to explore how breeding program structure affects the germplasm’s innate ability to create new genotypes with adaptation to environmental change and the balancing of the trade-off between optimal adaptation to the current TPE (i.e., short-term genetic gain) with adaptation to potential future TPEs (i.e., long-term genetic gain). This is done specifically in the context of hybrid varieties, which, since their emergence over a century ago (Shull 1908), have become the dominant variety type globally in most field and vegetable crops (Duvick 1999; Silva Dias 2010) and are credited as a significant contributor to global food security.

We have previously argued for the importance of considering the genetic complexity arising from highly interactive gene networks when studying the long-term behavior of selection programs in plant breeding (Cooper *et al*. 2005; Technow *et al*. 2021). One of the major features of complex genetic systems is that gene effects, and derivatives thereof, such as General and Specific combining ability (GCA and SCA), key measures in hybrid breeding (Sprague and Tatum 1942), are non-stationary and are dependent on the genetic background and population context in which they are observed (Wade 2002; Cooper *et al*. 2005; Huang and Mackay 2016; Powell *et al*. 2021; Milocco and Salazar-Ciudad 2022). This is even more obvious when considering the added complexity of environmental interactions. A trivial example is flowering time, where alleles conveying early flowering are beneficial in an early maturity environment but detrimental in a late maturity environ-ment, and vice versa (Li *et al*. 2018). Another such example are disease resistance genes, which are neutral or potentially even detrimental in the absence of the target pathogen (Burdon and Thrall 2003; Brown and Rant 2013). Therefore, we argue that it is necessary to consider the biological complexity resulting from interactions between genes and among genes and the environment to model and predict adaptation to long-term environmental changes (Podlich and Cooper 1999; Cooper and Podlich 2002). In this study, we build on the modeling framework previously developed by us (Technow *et al*. 2021) to investigate the implications of different breeding program structures on the balance between short and long-term genetic gain in a changing TPE. We do this on the basis of the graph theoretical *E(NK)* model of trait genetic architecture (Cooper and Podlich 2002), which extends Kauffman’s *NK* model (Kauffman 1993) with an environmental dimension. Our objective, as in our previous study, is not necessarily to identify and recommend superior strategies for implementation in practice. Rather our aim is to propose a framework to investigate and demonstrate the short and long-term behavior of these strategies in the context of the interplay between genetic and environmental complexity and change.

## Material and Methods

### Model of genetic complexity and environmental change

The *E(NK)* model developed by Cooper and Podlich (2002) on the principles of Kauffman’s *NK* model (Kauffman 1993) will form the basis of the simulations. Here *E* is representing the different ETs of the TPE, *N* (nodes in a graph) the number of genes for trait(s), as in the infinitesimal model, and *K* (edges in a graph) the average level of interactions among the *N* genes influencing the trait(s). Each *E(NK)* model comprises of a set of *NK* sub-models, one for each ET. Varying the number of ET, the degree of similarity between the *NK* sub-models, as well as the number of genes *N* involved in them and their degree of interaction *K*, creates a series of tunable models with differing environmental and genetic dimensionality and complexity. The use of the graph theoretical *E(NK)* framework for representing genetic architecture and its interaction with the environment was motivated by decades of research uncovering the biological networks underlying adaptation and output trait formation in globally important crops such as maize (e.g., Guo *et al*. 2014; Studer *et al*. 2017; Simmons *et al*. 2021; Tomura *et al*. 2025), rice (e.g., Wilkins *et al*. 2016) and wheat (e.g., Harrington *et al*. 2020).

#### *E(NK)* model implementation

We will first describe the *NK* sub-models and then how these were integrated into the *E(NK)* ensemble. The *NK* sub-models were implemented as in Technow *et al*. (2021), following the generalized approach described by Altenberg (1994), with the necessary adaptations for diploid genomes. Briefly, the complex trait is described as sum of a set of *fitness components*, normalized by dividing by *N* out of convention (Csaszar 2018). The values of the fitness components are distributed uniformly between 0 and 1 and are calculated as *random functions* of the genotypes at *K* interacting genes drawn at random from all *N* genes. The random functions were derived from the *ran4* pseudo-random number generator (Press *et al*. 1992). As in Technow *et al*. (2021), the number of fitness components and the number of genes were both set to *N* = 500. A number that reflects the very large number of genes and metabolites that are involved in development, plant architecture and biomass metabolism of complex organisms such as maize (e.g., Saha *et al*. 2011; Powell *et al*. 2022) while still remaining computationally tractable. Genes were biallelic and the simulated organism diploid. The range of the complexity parameter *K* considered was from 1 to 15 in steps of 1. For values of *K* > 1 we allowed for some variation in the number of interacting genes by sampling the gene number of each component from a Poisson distribution with rate parameter equal to *K* and then truncating the sampled values to fall between 1 and 15. At *K* = 1 all fitness components were controlled strictly by a single gene. For *K* > 1, genes were assigned at random to the fitness components, and individual genes consequently were typically involved in multiple fitness components, i.e., they acted pleiotropically. For *K* = 1, each of the 500 genes was assigned to exactly one fitness component at random and the values of heterozygous genotypes set to be mid-way between the alternate homozygous genotypes. Thus, at *K* = 1 genes acted strictly additively and without any intra- or inter-gene interactions. Because the expected value of the maximum of two samples from a Uniform distribution between 0 and 1 is 2/3, the expected maximum attainable fitness at *K* = 1 is also 2/3. The complexity of the resulting genetic space was quantified in our previous work (Technow *et al*. 2021).

The complete *E(NK)* model was comprised of 12 *NK* sub-models representing 12 discreet ET. This number was again a compromise between the large number of possible combinations of environmental factors (including crop management and pathogen and pest pressure) that can give rise to differential adaptation requirements, the typical number of ETs defined and targeted by a breeding program (e.g., Löffler *et al*. 2005), and computational tractability. The ET were assigned numbers from 1 to 12 on an ordinal scale. We will henceforth refer to the number distance between these numbers as “lag” (e.g., ET3 and ET6 have a lag of 3). The *NK* sub-models underlying each ET were created in such a way that the genetic correlation between the genotypic values in different ET, a common measure of the magnitude of GxE (Falconer 1952; Cooper and DeLacy 1994), decreased with increasing ET lag. To achieve this, 75% of the fitness components of each *NK* sub-model were retained unchanged from the sub-model of the previous ET and 25% were assembled anew as described above. Being the first in the series, the sub-model for ET1 was assembled completely from scratch. Thus, ETs with a lag of 1 will have 75% fitness components in common, ETs with a lag of 2 about 50% and so on. To directly quantify the resulting GxE correlation, 250 genotypes were generated at random and their fitness values calculated for each ET. The resulting correlation matrix was then summarized by averaging pair-wise ET correlations with the same lag. This process was repeated 300 times, each time using a new *E(NK)* model and 250 newly created genotypes. This was done at the intermediate value of *K* = 6, though preliminary analysis showed that results were consistent across all values of *K* (results not shown). The average GxE correlation by ET lag is shown in Figure 1(A). The genetic correlation between directly neighboring ET (lag = 1) was about 0.75, it dropped below 0.25 at around a lag of 5 and ET with a lag above 7 were virtually uncorrelated.

**Figure 1.**
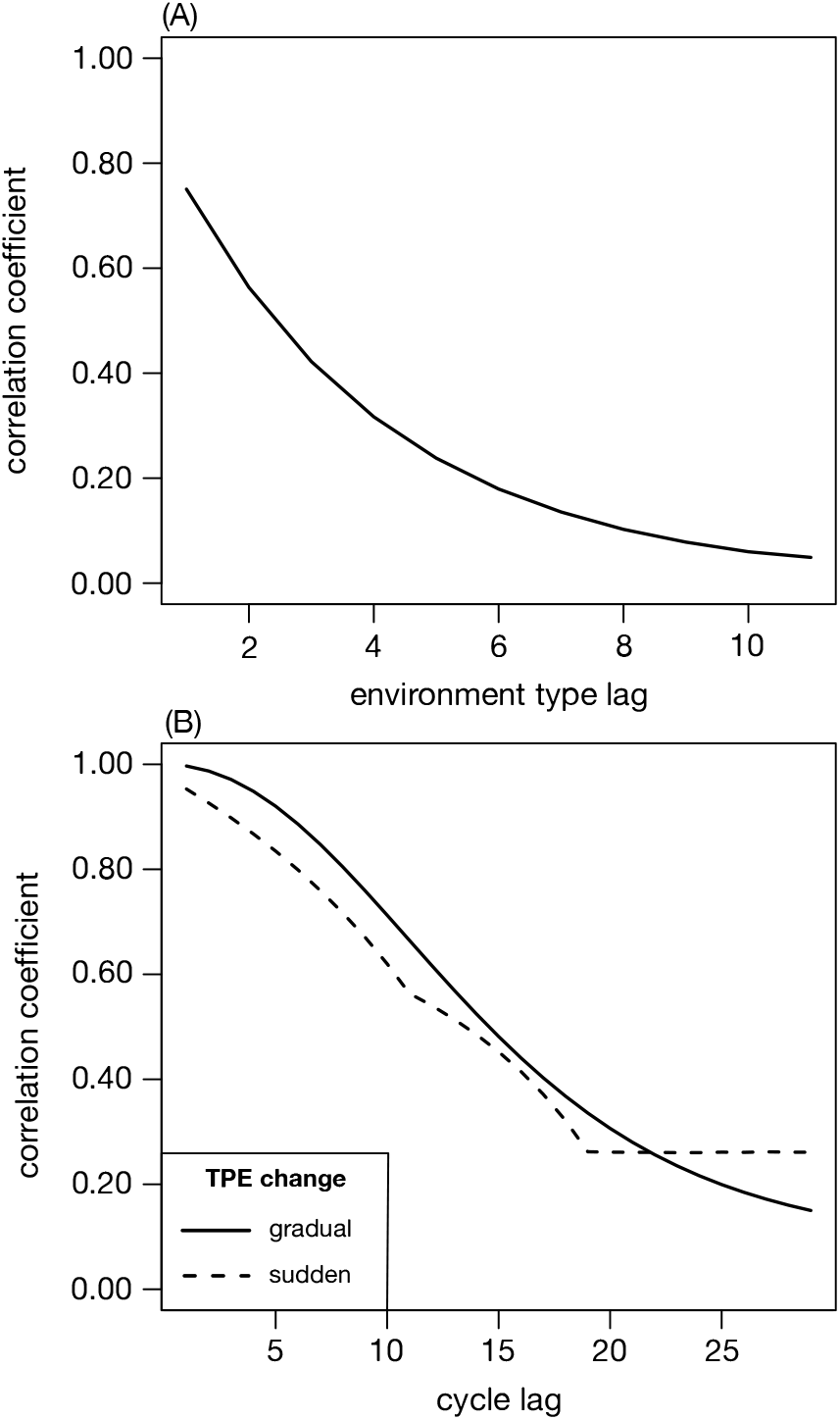
(A) Average correlations between the environment type (ET) specific performance as a function of ET lag. (B) Average correlations between across TPE performance lagged across cycles.

#### TPE change scenarios

A TPE can be described as a frequency distribution of ET (Cooper and Podlich 2002). Environmental change can thus be modeled as a change in those frequencies. Because the ET were modeled to be on an ordinal scale, this can conveniently be done by shifting the center (i.e., the area of highest probability weight) of the frequency distribution. We considered two environmental change scenarios taking place over 30 breeding cycles (1) *gradual* change and (2) *sudden* change. Under gradual change, the center of the ET distribution shifted gradually from ET1 in the first breeding cycle to ET12 in the last cycle (Figure 2). The most prominent example of this scenario are the incremental environmental changes brought about by climate change, such as the slow but steady rise in temperature and CO_2_ levels (Cooper *et al*. 2021). In the sudden change scenario, the weight of the frequency distribution remained centered over ET 1 to 6 for 20 breeding cycles and then shifted abruptly to being centered over ETs 7 to 12 for the remaining ones. This scenario represents the emergence of new pathogens and pests (McCann 2020), irrigation resources like the threatened Ogallala Aquifer in the western US corn belt reaching a critical depletion level (Basso *et al*. 2013) or enactment of new agricultural policies, like those regulating the application of nitrogen fertilizer (Kanter *et al*. 2020).

**Figure 2.**
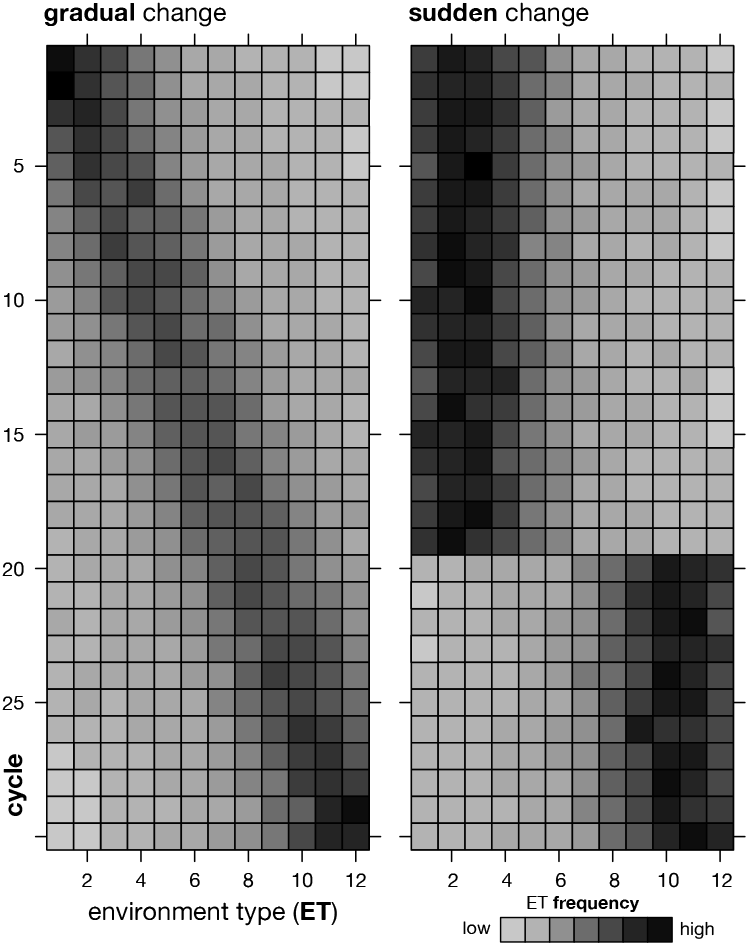
Visualization of change in ET frequency distribution for representative examples of the gradual (left) and sudden (right) TPE change scenarios

The actual ET frequencies were sampled from Dirichlet distributions with concentration parameters chosen such that the center of the ET frequency distribution followed the general TPE change pattern outlined above. Sampling rather than setting these frequencies directly mirrors random year over year variability, in e.g., weather patterns. Because the support of the Dirichlet distribution is strictly positive, all ET had non-zero frequencies. The majority of the frequency distribution, however, was concentrated over a few ET around the current center of the distribution. This can be quantified with the “effective number” of ET, calculated as 1/***b***′***b***, where ***b*** is the vector of ET frequencies. This corresponds to the “Inverse Simpson Index” concentration measure used in e.g., ecology. The average values were 5.9 and 5.4. for the gradual and sudden TPE change scenarios, respectively. Thus, less than half of all 12 ET had an effective presence in the TPE at any given point in time.

True performance values of individuals in a given TPE were calculated as the average fitness values across the separate ET, weighted by their frequencies in the current TPE. The GxE correlation between TPE performance values across different cycles was calculated similarly as the GxE correlation between ET, i.e., from the true performance values of 250 randomly generated genotypes across all 30 cycles at a complexity level of *K* = 6. This was also repeated 300 times with newly generated genotypes and *E(NK)* models. Results are shown in Figure 1B. Consistent with expectations, the GxE correlation gradually decreased with increasing cycle lag in the gradual TPE change scenario. The lag curve for the sudden change scenario is included for completeness but it is less meaningful as a descriptor of the GxE correlation between cycles in this scenario. Here, the only directional change in the TPE occurred at cycle 20. Consequently, the GxE correlation between the TPEs in cycles 1–20 and 21–30 was close to one and would have been equal to one if not for the small random frequency differences imposed. The GxE correlation between cycles with different TPEs (e.g., between cycle 8 and 23) was 0.266 and similar in magnitude to the GxE correlations between the largest cycle lags in the gradual scenario.

Wright (1932) introduced the metaphor of a *genetic landscape* as a conceptualization of high-dimensional genetic complexity. Naturally it has also been found useful for developing an intuition for the ruggedness of different *NK* models (e.g., Kauffman and Weinberger 1989). While it should be emphasized that as a metaphor it should not be taken literally, we too find it helpful in making accessible the rather abstract concepts surrounding the discussion of the properties of *E(NK)* models. Using the terminology of Technow *et al*. (2021), the landscape at *K* = 1, where genes act strictly additive without any inter- or intra-gene interactions, can be visualized as *Mount Fuji*, i.e., an isolated peak with monotonous incline to the top (Figure 3). At intermediate levels of *K*, the genetic landscape contains a cluster of peaks, like a mountainous region in an otherwise flat landscape, similar to the *European Alps*. Finally, at high values of *K*, the landscape can be thought of as a *Sea of Dunes*, i.e., an endless number of evenly distributed peaks of similar shape and size. The different *E(NK)* sub-models representing the ET would then correspond to a series of different landscapes with more or less similar features. And TPE change in particular could be thought of as landscape change, with old peaks flattening and new ones appearing as a result of geological forces.

**Figure 3.**
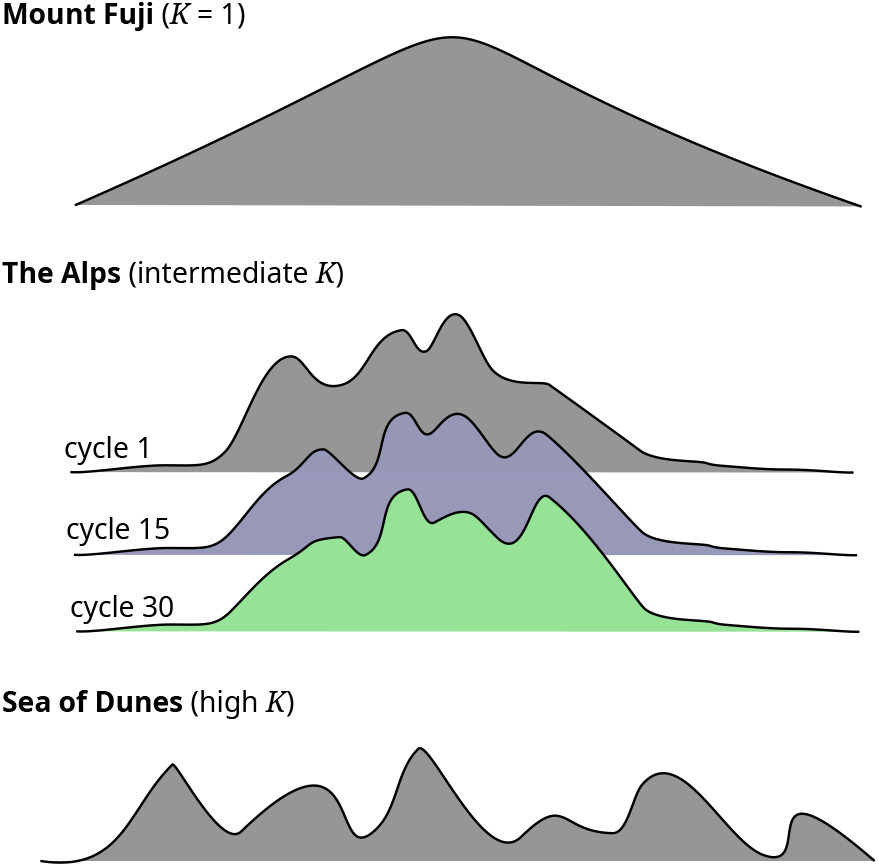
Schematic visualization of genetic landscapes corresponding to different values of complexity parameter *K*. The different colors in the ‘The Alps’ schematic represent the changing genetic landscape resulting from environmental change over cycles. Figure modified with permission from Figure 2 of Technow *et al*. (2021)

#### Genome definition

The simulated genome consisted of 10 diploid chromosome, each of 1 Morgan in length and with the *N* = 500 genes distributed evenly across them. Meiosis and recombination was simulated according to the properties of the Haldane mapping function with the R package *hypred* (Technow 2013), which is available from the supplement of Technow and Gerke (2017).

### Simulation of hybrid breeding process

The simulation process followed closely the one in our previous work (Technow *et al*. 2021), which was designed to reflect the features of long term hybrid breeding operations observed in practice (Duvick *et al*. 2004; Mikel and Dudley 2006). It is visualized in Figure 4 for the most general *distributed* program structure, of which the others are special cases (more details on this and the other structures will be given below). On the outset, a base population comprising 1,000 inbred lines was generated stochastically with the approach described by Montana (2005). This was done in such a way that the expected LD between two loci *t* Morgan apart equaled *r*^2^ = 0.5 · 2^−*t*/0.1^ and all minor allele frequencies were distributed uniformly between 0.35 and 0.50. This base populations was then separated at random into two equally sized heterotic groups (arbitrarily labeled ‘1’ and ‘2’) and, depending on the scenario, further into sub-populations within those. Sub-heterotic patterns where then formed by pairing one sub-population from one heterotic group with one sub-population from the other group. We will subsequently refer to those pairs also as “breeding programs” or simply as “programs”. Hybrids were produced strictly by crossing lines across heterotic groups while breeding crosses to generate the next generation of recombinant lines where done strictly within the designated heterotic groups. However, while hybrid crosses were done only within a heterotic pattern (i.e., program), breeding crosses could be done done within and among sub-populations, depending on the scenario (Figure 4B).

**Figure 4.**
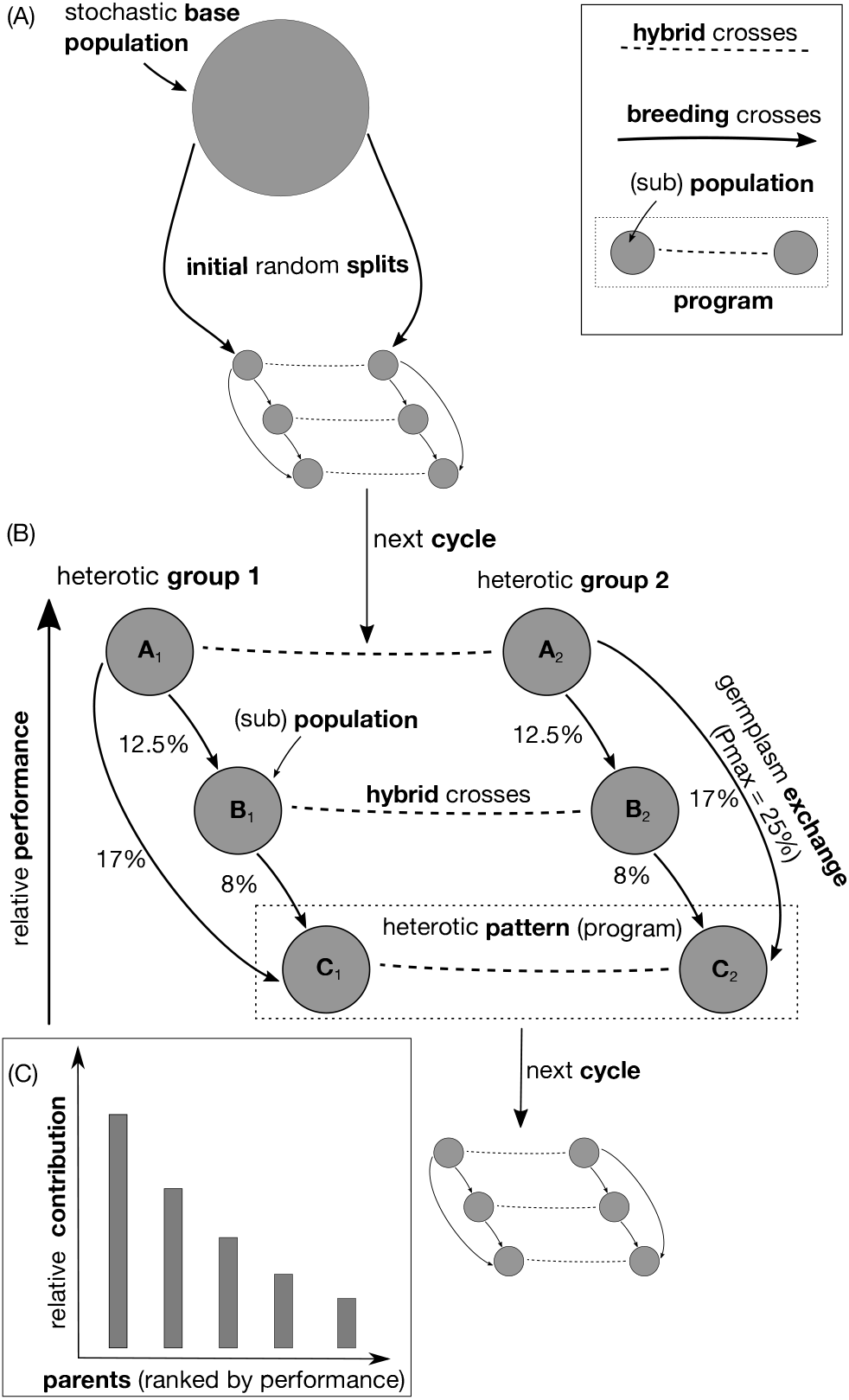
Schematic visualization of simulated hybrid breeding process using the distributed program structure as example. (A) Stochastic creation of base population and separation into heterotic groups and sub-populations; (B) detailed snapshot view of the processes happening within each cycle; (C) relationship between the performance of selected breeding parents and their relative contribution to the next generation. The programs were ranked based on the average performance of their experimental hybrids in a given cycle. The percentage values next to the arrows connecting programs indicate what percent of the breeding crosses are conducted with lines from higher ranked program. The lower the performance rank of a program, the higher this percentage, with lowest ranked program receiving the maximum given by Pmax and the highest ranked program not conducting any crosses with outside line. The figure was reproduced with permission from Technow *et al*. (2021).

#### Evaluation of genetic performance

The generalized combining ability (GCA) of each line, the primary selection metric used in hybrid breeding (Reif *et al*. 2005), was evaluated with an incomplete, reciprocal mating design (Melchinger *et al*. 1987; Seye *et al*. 2020) by performing crosses with five random lines from the sub-population of the opposite heterotic group of the same program. The performances of the resulting testcross hybrids were then obtained with the *E(NK)* model as previously described and averaged. A normally distributed noise variable with zero mean was then added to reflect residual noise. The variance of this noise variable was chosen in such a way that the resulting GCA values had a heritability of 0.75 on an entry mean basis, a value achieveable for traits like yield in multi-environment, multi-testcross trials (Schopp *et al*. 2015).

The so obtained “observed” GCA values were then also used to predict the performance of all possible inter-group hybrids within the program by summing the GCA values of the corresponding parent lines (Reif *et al*. 2005). The top hybrids were then selected based on these GCA predictions and their true performance determined according to the *E(NK)* model. The size of this selected class, which represents the set of advanced experimental hybrids considered for commercial release, depended on the scenario and more details will be given below. The true performances of these experimental hybrids were then averaged and this average used to quantify the overall performance of the program in a given cycle for the purpose of ranking the programs for determining the proportion of breeding crosses with lines from other programs, as will also be described below in more detail. Commercial breeding programs typically release only very few hybrids from each cycle to farmers. The performance of the top experimental hybrid across all programs, i.e., the one that would be targeted for commercial release in practice, was consequently defined as the practically relevant performance metric for the whole breeding operation in that cycle and also used as a metric of genetic gain.

#### Selection of breeding crosses

To determine which lines contributed to the next generation through breeding crosses and by how much, they were assigned a usage probability that was the product of an individual and population level relative contribution value. The individual level contribution value was a function of the rank of a line within its sub-population and determined as follows. First, the lines were ranked according to their observed GCA values (Figure 4C). Low ranked lines received a contribution value of zero, which excluded them as potential breeding cross parents. The contribution values of the top ranked lines within each sub-population were then drawn from a Dirichlet distribution. The concentration parameters of this distribution were chosen in such a way that the relative contributions halved with every 1/5th rank quantile. Thus, with, e.g., 25 lines selected as parents, the highest performing line contributed approximately twice as much to the next generation as the 5th ranked line and four times as much as the 10th ranked line. How many lines were selected depended on the scenario and will be detailed when describing the alternative breeding program structures. These individual contributions thus correspond to the *disproportional* contribution scenario as defined in Technow *et al*. (2021), and reflect the practical reality observed in long-running commercial breeding operations (Rasmusson and Phillips 1997; White *et al*. 2020).

The population level contribution values regulate how much the lines of one program contribute to the breeding crosses of another. The example visualized in Figure 4B will be used for describing how they were derived. In each cycle, the programs (labeled ‘A’, ‘B’ and ‘C’, with subscript 1 or 2 indicating the heterotic group) are ranked according to the average performance of their experimental hybrids, as described above. Generally, the lower the performance rank of a program, the higher the proportion of breeding crosses with lines from other programs, from zero for the highest ranked program (A), which consequently does not use any outside lines in breeding crosses that cycle, to a value of Pmax for the lowest ranked program (C) and equidistantly spaced values for the program(s) with intermediate ranks (B). Furthermore, exchange of germplasm between programs happened strictly in the direction from higher to lower ranked programs. In the example as well as throughout this study, Pmax was 25%. Thus in the example, the highest ranked program A performs no crosses with external lines, the intermediately ranked program B uses lines from other programs in 12.5% of its new crosses and the lowest ranked program C uses lines from programs A and B in Pmax = 25% of its crosses. The exact proportion derived from each of the higher ranked programs was proportional to how much their performance differed relatively. If, as in the example, the performance difference between programs C and A is twice as large as that between C and B, lines from program A will contribute twice as much to program C than lines from program B. Specifically in this example 17% from A and 8% from B for at total of 25%. Because the ranking and relative performances of the programs can change over cycles, those relative contributions are derived anew each time as well. As stated, use of lines for breeding crosses, within and across sub-populations, happened strictly within their respective heterotic groups, e.g,. sub-population C_1_ could receive lines only from A_1_ and B_1_. This scheme reflects that in practice, successful programs have little incentive to use outside material for improving their genetic base while less successful programs will do so at increasing rates in order to close the observed performance gaps. To arrive at the final usage probabilities, the individual and population level contributions were then multiplied with each other. Then the breeding crosses were determined by sampling with replacement the lines proportionally to their usage probabilities. While this could result in multiple instances of the same cross (i.e., between high performing lines), selfings were excluded. From each of these crosses one recombinant line was obtained through three generations of single seed descent selfing, followed by a final doubled haploidy generation to remove residual homozygousity. These new recombinants then formed the next breeding cycle, fully replacing the previous generation. Generations were thus discrete and a line could be used as a breeding cross parent in only one cycle. The breeding simulation was run for 30 cycles and 350 independent repetitions were conducted for each of the scenarios studied. Unless specified otherwise, all results presented are the averages over these 350 replications.

### Alternative breeding program structures

Following Technow *et al*. (2021), we distinguished and explored three main alternative scenarios for how breeders might choose to structure their breeding programs. The already mentioned *distributed* structure (Figure 4) is characterized by the presence of multiple smaller programs that interact with each other through the exchange of genetic material in the form of breeding cross parents, as described in the previous section. In the *centralized* structure only a single, large program exists, with the only separation being into heterotic groups, which is fundamental for hybrid breeding (Melchinger and Gumber 1998). The final structure considered in our previous work was the *isolated* structure (*iso*), which resembled the distributed structure, except that no exchange of genetic material across sub-populations took place (i.e., Pmax = 0%). Here we add two further structures where genetic exchange took place, but only sporadically. These could thus be considered as intermediates of the distributed and iso structures. In the *iso7* structure, exchange happened every 7th cycle and in the *iso21* structure only once in cycle 21. In the cycles in which exchange took place, it happened exactly as already described for the distributed structure. Note that in iso21, the cycle in which exchange took place immediately followed the TPE shift under the sudden TPE change scenario, as a dramatic environmental change might prompt breeders to break their isolation temporarily and seek out genetic material from programs that navigated the shift better. Before the exchange cycle, the iso and iso21 structures operate identically and no differences between them should be expected.

The distributed and isolated structures comprised five programs each with one sub-population of size 100 per heterotic group. In every cycle, 25 lines were selected from each sub-population as potential breeding cross parents. The number of experimental hybrids selected per program was 25. In the centralized structure the number of lines per heterotic group was 500, with 125 selected as parents. Here, the number of experimental hybrids was 125. The total number of lines and selected breeding parents per heterotic group as well as experimental hybrids was with 500, 125 and 125, respectively, the same across all structures. Thus, the alternative structures can be considered as using approximately the same amount of resources in this regard. Because of the uneven use of parental lines in crosses and the possibility of them contributing to multiple programs, we quantified for each cycle the effective number of breeding cross parents contributing to the next generation with the previously defined Inverse Simpson Index, with ***b*** now being the vector of relative contributions (Boichard *et al*. 1997). This was done across the sub-populations within the arbitrarily chosen heterotic group 1. Averaged across cycles, the resulting values were 61.2 (distributed), 62.0 (centralized), 64.6 (iso7), 65.0 (iso21) and 65.1 (iso).

### Recorded metrics

Several metrics were used to study the short and long-term behavior of the simulated breeding system.

1. The performance value of the top hybrid identified in a given cycle from each structure was defined as the performance of this structure in that cycle.
2. The proportion of GCA to total genetic variance (%GCA) is a key metric describing the amount of additive genetic variation exploitable for heritable genetic improvements, as well as for identification of superior hybrid combinations in a given cycle. It was calculated as described in Technow *et al*. (2021), except that we used the *MCMCglmm* R package (Hadfield 2010) to estimate variance components. %GCA was estimated separately for each program, for the structures where there were multiple, and then averaged to arrive at a single value per cycle.
3. The correlation between GCA effects (GCA correlation) was used as a measure of transferability of genetic effects across programs. It was evaluated by performing testcrosses with partners across sub-heterotic patterns. For example, the lines from A_1_ were testcrossed with lines from B_2_ and the so obtained GCA values correlated with those from the regular testcrosses with A_2_ (Figure 4). For computational reasons, this was done for only one sub-heterotic pattern and only every third cycle.
4. The global Fst statistic was used to quantify genetic differentiation between sub-populations in the distributed and isolated structures. Fst values between populations were calculated separately for each heterotic group and then averaged, i.e., they quantify differentiation within a heterotic group. The Fst statistics were calculated with function fs.dosage from the *hierfstat* package in R (Goudet and Jombart 2022).
5. Following Technow *et al*. (2021), we used the proportion of loci with a minor alleles frequency below 5% (*U*_*W*_ ) as a measure of allelic diversity. This measure also quantifies the thickness of the extreme tail of the allele frequency distribution. The higher it is, the more the distribution is ‘*U-shaped*’, which was identified as a key determinant for the amount of additive genetic variation observed in natural populations (Hill *et al*. 2008). This calculation was done separately for each sub-population and then averaged to arrive at a single value per cycle for each.
6. The effective population size (*N*_*e*_), calculated according to the method described in Corbin *et al*. (2012) for estimating constant effective population size, was used as a holistic measure of the diversity and coverage of genetic space. It was calculated across the sub-populations within each heterotic group and then averaged.
7. The ranking of sub-populations determines flow of genetic material and reflects how much each contributes to overall, commercially relevant, genetic gain (measured as performance of top performing hybrid in a given cycle). We therefore defined a metric called *top-rank stability* which represents the probability that the top-ranked program in cycle *n*−1 is also the top-ranked program in cycle *n*. The value for cycle *n* was obtained as the proportion of the replications of the simulation in which the top-ranked program in cycle *n* was the same as the one in cycle *n* − 1. To aid interpretability, the resulting curve across cycles was smoothed using loess.

All computations were conducted in the R environment for statistical computing (R Core Team 2021).

## Results

### Absolute performance over time

For brevity, results will only be shown for *K* values of 1, 8 and 15, representing genetic additivity, and intermediate and high genetic complexity, respectively.

#### Gradual TPE change

At *K* = 1, performance in all structures increased until about cycle 10 but then leveled off or even decreased, as particularly pronounced in the case of the iso structure (Figure 5). During intermediate cycles, the centralized structure had the highest performance, followed by the distributed and iso7 structure. A noteworthy exception to the general trend of stalling or declining performance in later cycles was structure iso21, for which performance increased again noticeably after the germplasm exchange event in cycle 21. At *K* = 8 the isolated structures reached a higher performance level quicker and were superior during the first 20 cycles, after which performance stalled and then dropped off towards the end. The distributed structure followed the isolated structures closely and replaced them as the superior structure by cycle 25 as performance kept increasing throughout. The centralized structure saw noticeable cycle over cycle genetic performance improvements only after cycle 15, after which performance improved at a faster rate than for the other structures. At *K* = 15 none of the structures showed any performance improvement until cycle 10. From there on, performance increased for the isolated structures, with a slowing rate towards the end. The iso21 structure very closely followed the iso structure, as did iso7, albeit with a slightly lower rate of gain. Performance for the distributed structure started to make improvements after cycle 15, for the centralized structure it stayed virtually flat throughout.

**Figure 5.**
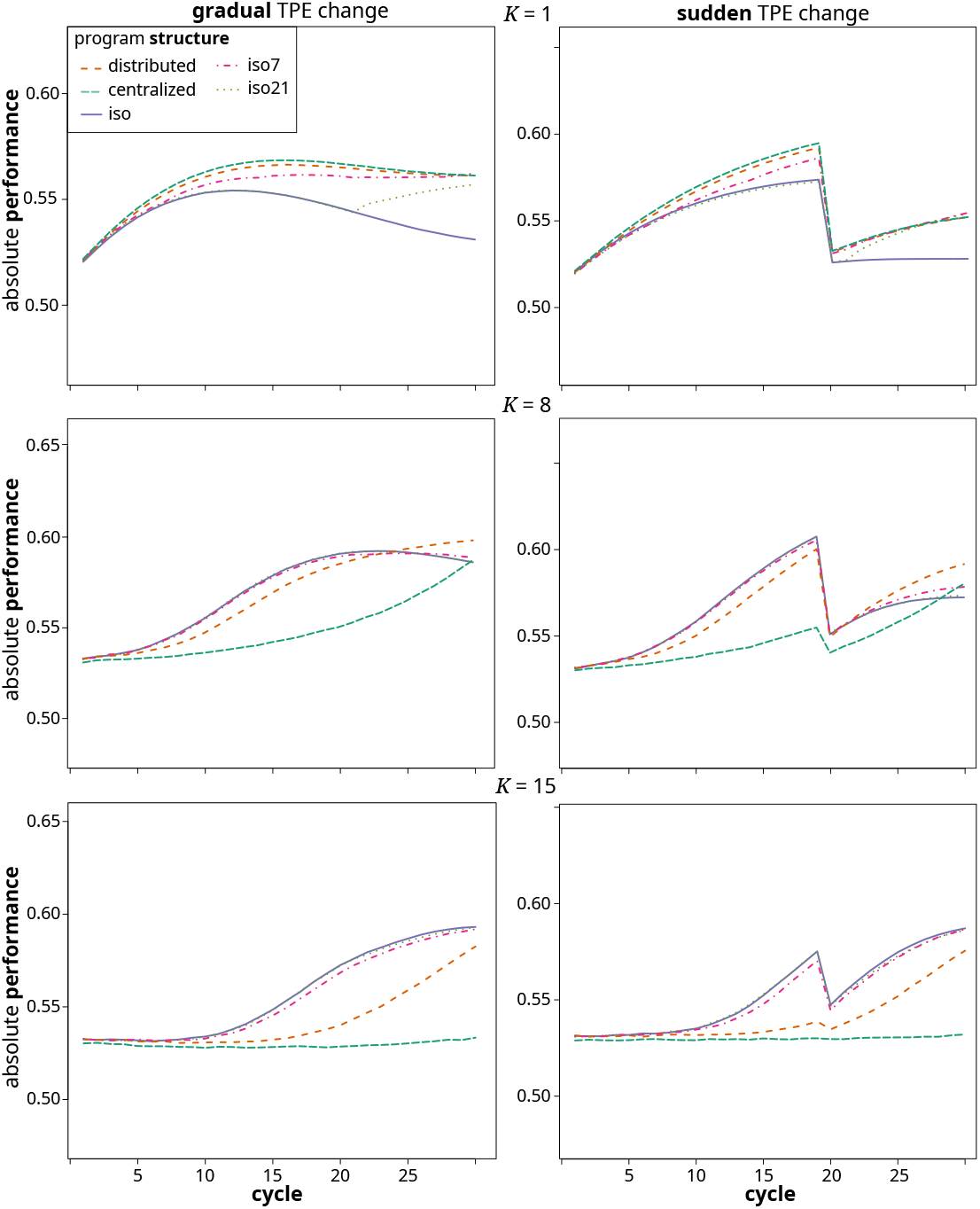
Absolute performance across cycles of the different breeding program structures in the gradual (left) and sudden (right) TPE change scenario at *K* levels of 1 (top), 8 (middle) and 15 (bottom).

#### Sudden TPE change

At *K* = 1, performance increased for all structures from cycle 1 on, with the centralized structure reaching the highest level until the TPE shift, followed by the distributed and iso7 structures. Performance increase in the iso and iso21 structures was lowest. After the TPE change at the end of cycle 20, performance collapsed dramatically to close to the initial value for all structures. After this, it remained flat in the iso structure, whereas it started to recover again in all others and in a similar way. At *K* = 8 and 15, a similar pattern as in the gradual scenario was observed, except that like for *K* = 1, performance collapsed after the TPE change and then started to recover. In contrast to *K* = 1, this recovery was also observed for the iso structure.

### Relative ranking of program structures over time

The relative performance of the different structures was visualized for all combinations at each combination of TPE scenario, *K* and cycle in Figure 6. This figure represents the relative ranking of each structure with colors scaled to be situated linearly between the lowest and highest performing structure.

**Figure 6.**
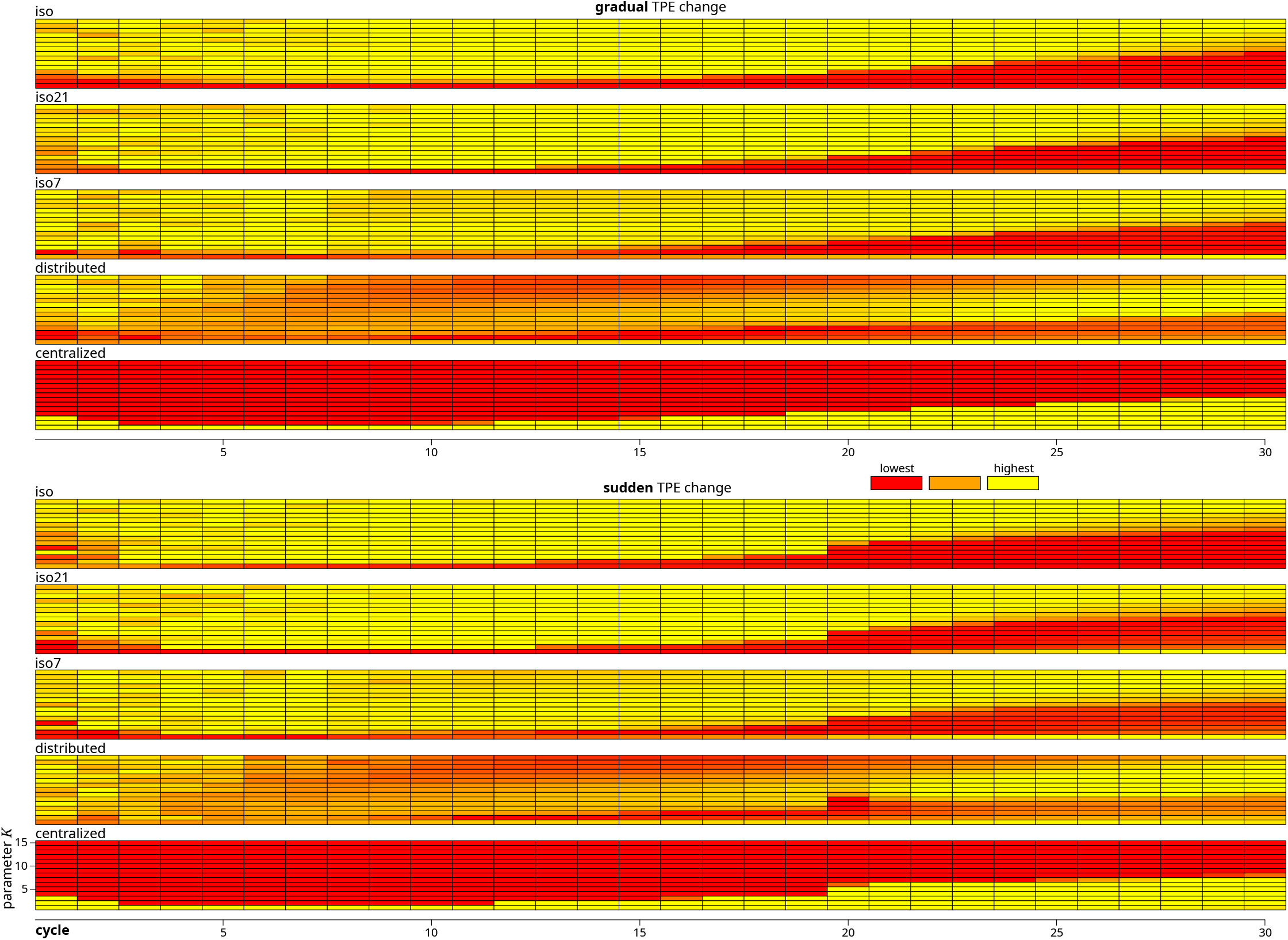
Relative performance ranking of structures across cycles, *K* levels and TPE scenarios. The brighter the color, the higher the relative performance of the structure compared to the others. Color values were linearly scaled to the range between the highest and lowest performing structure within this cycle, *K* and TPE combination. The values represented are the average performances across the replications of the simulation.

In both TPE change scenarios, the isolated structures (iso, iso7 and iso21) generally were the superior structures during the first 15 cycles, except under *K* values below 3 in the very first cycles and throughout for *K* = 1, where the centralized structure was superior . The iso7 structure generally ranked very similarly to the iso structure, with the notable exception being *K* = 1, where the iso structure was the lowest ranking throughout, while iso7 maintained an intermediate rank. Similarly for iso21 after the the exchange cycle. During the later cycles the isolated structures generally were superior only at higher levels of *K*. For intermediate values of *K* the distributed structure was superior and for lower levels of *K* the centralized structure. The distributed structure throughout the cycles and across levels of *K* when not superior maintained at least an intermediate rank.

### Quantitative Genetic Parameters

The various quantitative and population genetic parameters used to measure system behavior showed the same general pattern across both TPE change scenarios. For sake of brevity we will therefore illustrate these only for the gradual scenario. We will also highlight results only for *K* levels of 1, 8 and 15.

**%GCA** For *K* = 1, %GCA was equal to 100% for all structures, as expected (Figure 7). At *K* = 8 %GCA started from just above zero and then increased from there. This happened fastest in the iso and iso21 scenario, which reached above 50% before cycle 15 and close to 100% by the final cycle. The iso7 structure followed slightly behind the other two. It also showed a sudden small increase after the second and third germplasm exchange cycle and a sudden decrease after the final exchange cycle 28. iso21 also showed a small increase after its exchange cycle. All of these were short lived and the effect dissipated the following cycle. %GCA increased slower in the distributed structure, which reached 50% only by cycle 15 and just above 80% by the final cycle. The increase was slowest in the centralized structure, which had barely reached 40% by the final cycle. These trends were similar but more pronounced at *K* = 15, where the iso7 structure now fell noticeably behind the iso structure, the distributed structure reached 50% only around cycle 25 and the centralized structure remained at close to zero throughout.

**Figure 7.**
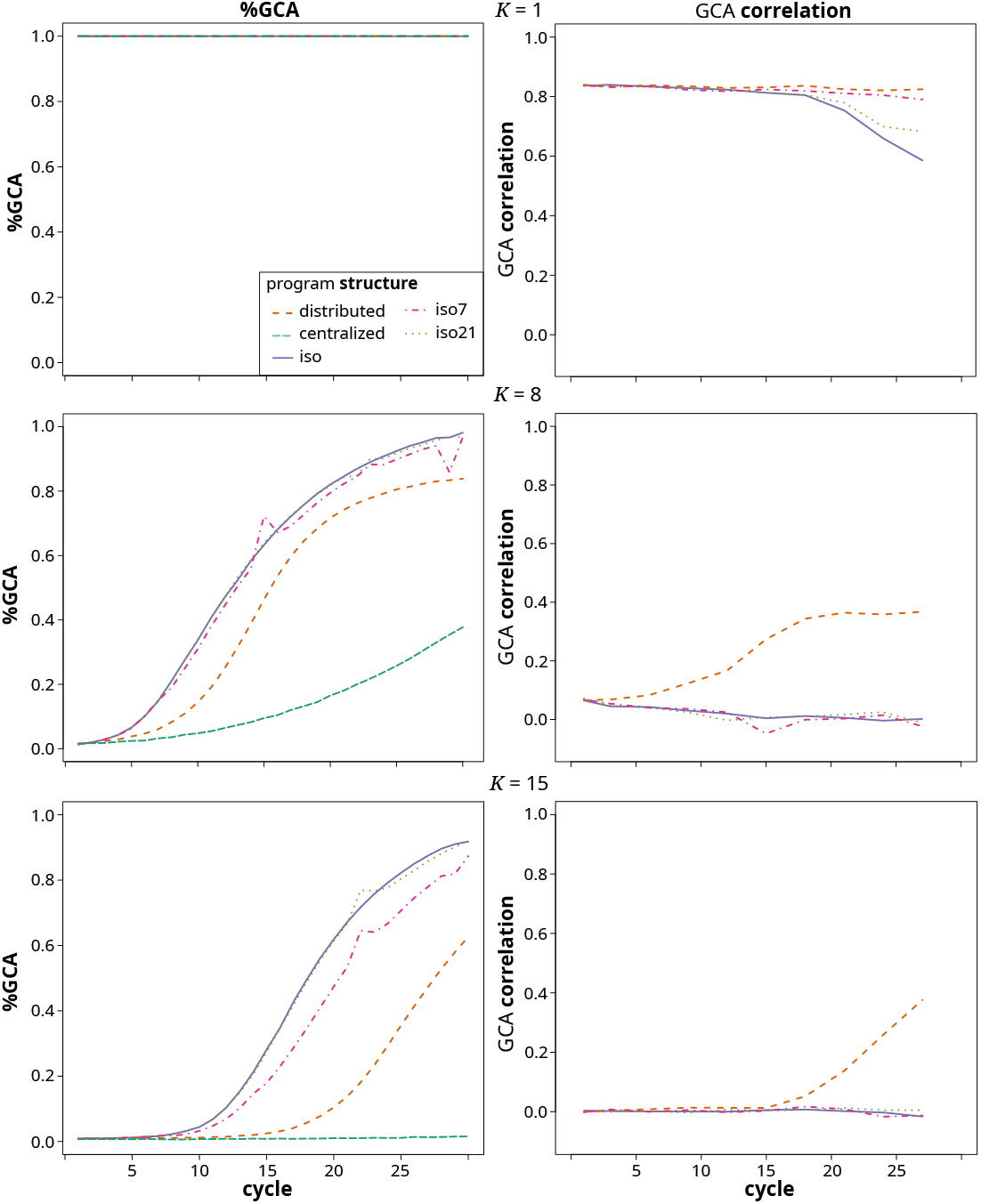
%GCA (left) and GCA correlation (right) across cycles for the different breeding program structures in the gradual TPE change scenario at *K* values of 1 (top), 8 (middle) and 15 (bottom). The values represented are the averages across the replications of the simulation. Note that the GCA correlation does not apply to the centralized structure.

#### GCA correlation

At *K* = 1, the GCA correlation throughout remained at close to 0.85 for the distributed and iso7 structures (Figure 7). For the iso and iso21 structures it started falling after cycle 15, with a stronger decline in the iso than the iso21 structure. At *K* = 8, the correlation was below 0.1 initially and declined further towards zero for the iso, iso7 and iso21 structures. In case of the distributed structure, the correlation increased until it reached a plateau of just below 0.4 after cycle 20. At *K* = 15 the correlation remained at zero for all structures, except the distributed structure, where it started to increase from cycle 15 onward and reached close to 0.4 by cycle 27.

#### Fst

At *K* = 1, Fst among sub-populations in the distributed structure quickly increased to a value of 0.17 and then remained there for the remainder (Figure 8). In the iso structure it quickly rose to above 0.6 by cycle 10 and ultimately reached 1.0 in the final cycles. The iso7 and iso21 structures shared the high rate of Fst increase. While Fst dropped after each exchange cycle, it rose again immediately thereafter. For the iso7 structure that led to Fst oscillating around a value of 0.5 from cycle 7 onward. At *K* = 8 Fst in the iso structure again rose quickly and reached 0.5 before cycle 10. The iso7 and iso21 structures again showed drops in Fst after germplasm exchanges. In contrast to *K* = 1, the Fst value prior to the exchange was regained after just two cycles. Fst in the distributed structure rose more slowly than in the other structures and remained at around 0.47 from cycle 23 onward. The trends at *K* = 15 were similar to those at *K* = 8, except that the Fst increased at a slower rate in all structures, values in iso7 were now noticeably lower than in iso and iso21 and in the distributed structure they did not reach a plateau and were lower overall.

**Figure 8.**
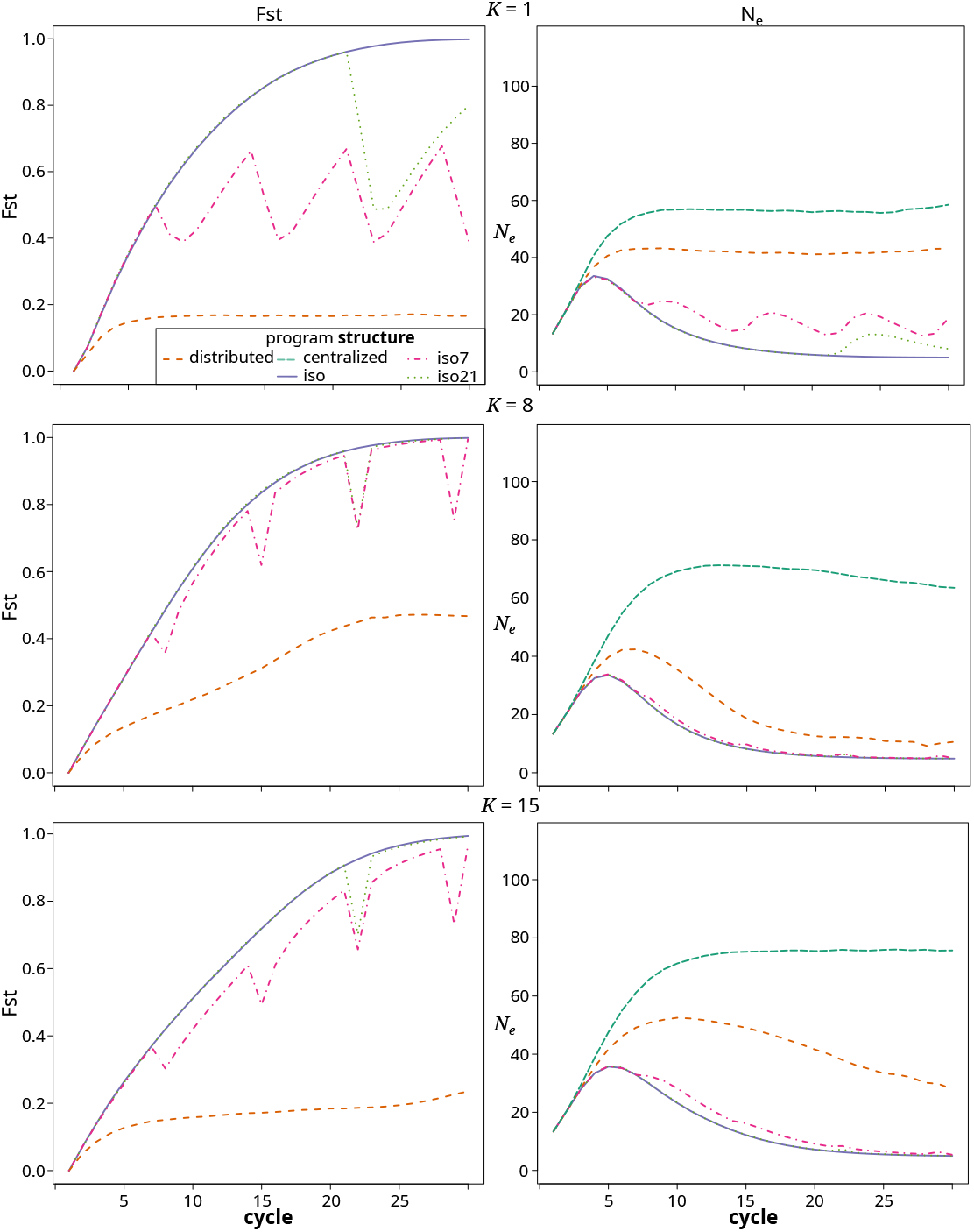
Fst (left) and *N*_*e*_ (right) across cycles for the different breeding program structures in the gradual TPE change scenario at *K* values of 1 (top), 8 (middle) and 15 (bottom). The values represented are the averages across the replications of the simulation. Note that the Fst among programs is not applicable to the centralized structure.

#### Effective population size

At *K* = 1, *N*_*e*_ across sub-populations in the centralized and distributed structures rose from the initial value and quickly stabilized at around 55 and 42, respectively (Figure 8). In the different iso, iso7 and iso21 structures, *N*_*e*_ rose initially until cycle 5 but then dropped again. In the iso and iso21 structures it converged towards a value of 5, which is the number of sub-populations within each heterotic group. After each exchange cycle in the iso7 and iso21 structures, *N*_*e*_ increased briefly but then dropped again. This led to an oscillating pattern around a value of 17 in iso7. At *K* = 8, the centralized structure again had the highest *N*_*e*_, which reached above 70 at the peak around cycle 10 before slowly decreasing again. In contrast to *K* = 1, *N*_*e*_ of the distributed structure initially increased to above 40 but then started falling again and ultimately stabilized at around 10. The pattern for the iso, iso7 and iso21 structures was similar to *K* = 1, except that the effect of the germplasm exchanges on *N*_*e*_ was barely noticeable. The pattern at *K* = 15 was similar to *K* = 8, except that the decrease in *N*_*e*_ after the early peak in the distributed and isolated structures was slower. In the distributed structure *N*_*e*_ reached above 50 at its peak and was still above 25 in the final cycle. In the three isolated structures, it nevertheless had declined towards 5 towards the end.

#### U-shapedness

The proportion of loci with MAF *<* 0.05 within sub-populations (*U*_*W*_ ) increased across cycles and program structures and levels of *K*. The increase was always strongest in the iso structure, where it reached a value of close to 100% in the final cycles. At *K* = 1 it increased with a similar overall rate and reached a similar final value in all other structures. The iso7 and iso21 structures showed a drop after each exchange cycle, followed by a renewed increase. At higher complexity levels, this drop after the germplasm exchanges was even more pronounced but also was reversed quickly in the following cycle. For the distributed and centralized structures the increase was slower at higher *K* values, strongly so for the latter, where *U*_*W*_ reached just about 20% by the final cycles.

#### Top-rank stability

At low *K*, the top rank stability was highest in the iso structure, where it increased over time to close to 100% and lowest in the distributed structure, where it remained constant at just below 40% (Figure 10). In the iso7 and iso21 structures, the value dropped after each exchange cycle, followed by a relatively rapid increase to the prior level. At intermediate and high values of *K*, it increased from around 20%, the value indicating randomness, to 75% and higher in all structures. The exception was the distributed structure at intermediate *K*, where it quickly rose to close to 100% by cycle 15 but then dropped again to 60%.

## Discussion

The existence of additive genetic variation, a prerequisite for achieving heritable selection gains, is a given only in the simplest gene to phenotype model of purely additive gene action. As a model of biology this seems inadequate on the basis of scientific advances in the field of biology demonstrating that quantitative traits are the product of complex interactions at the molecular, metabolic and physiological level (e.g., Carlborg and Haley 2004; Phillips 2008; Saha *et al*. 2011; Wilkins *et al*. 2016; Boyle *et al*. 2017; Fiévet *et al*. 2018; Vasseur *et al*. 2019). Infinitesimal and purely additive gene action also seems inadequate to explain why commercial plant breeding operations have managed make impressive levels of genetic gain, despite having structures that should not be conducive for this success under this model (Rasmusson and Phillips 1997; Duvick *et al*. 2004; Technow *et al*. 2021) or for the continued response in scientific long-term selection experiments (Dudley and *Technow et al. 11* Lambert 2010*). Under all other models of genetic complexity, heritable additive genetic variation is an emergent property of the non-linear gene networks underlying phenotypic variation that has to be “exposed” by constraining genetic variability in these networks (Wade 2002; Cooper et al*. 2005) through fixation or near fixation of genes and gene complexes (Hill *et al*. 2008). A process leading to the effective simplification and linearization of these networks.

With this background, we previously investigated how an increasing degree of genetic complexity affects the performance and properties of different hybrid breeding program structures (Technow *et al*. 2021). There we found that as genetic complexity increases, breeding program structures applying increasing constraints on the accessible genetic variability through population sub-division are required to make meaningful short- and long-term genetic gain. Here we explored this topic further by adding the dimension of environmental complexity, in the form of a gradually or suddenly changing TPE.

### Balance between short and long-term selection response

In Technow *et al*. (2021) we evaluated the relative superiority of the alternative structures in terms of their performance in the final cycle of the breeding simulations. The simulated scenario there was that of a static TPE, where the performance value in the final cycle is a reflection of how successful a structure was in traversing the static genetic landscape in search for higher and higher local optima or even the global one.

With a dynamic TPE, however, the genetic landscape changes as well (Figures 1, 2, 3). The local optima achieved in the final cycle are thus not indicative of the local optima identified in earlier cycles and hence how well and quickly a given breeding structure was able to react to historical shifts in the genetic landscape. For reference, even with deployment of all currently available breeding technologies like doubled haploids (Dwivedi *et al*. 2015), whole genome selection (Meuwissen *et al*. 2001) and evaluation in virtual MET and high throughput phenotying (Cooper *et al*. 2022; Voss-Fels *et al*. 2019), each breeding cycle, from initial cross to the point where the next generation of proven, elite lines are recycled, will require three to five calendar years (Heffner *et al*. 2010). The 30 cycles covered in this simulation thus easily span a whole century and with that the potential for massive shifts in the biotic and abiotic environment, management practices and market demands (Collins and Chenu 2021; Zhao *et al*. 2022; Shi *et al*. 2025).

We therefore here assessed the balance of short to medium and long-term selection response under a shifting genetic landscape, by evaluating whether the different structures were able to both maneuver and achieve genetic gain within the current genetic landscape while maintaining the ability to maneuver also in future, yet unknown landscapes.

In general, we found that earlier during the earlier cycles (1-10), the isolated structures generally delivered superior rates of performance improvement, followed by the distributed structure and with the centralized structure faring poorest. For the medium and long-term we found that the isolated structures required increasingly higher degrees of complexity to retain their superiority. As on the one hand their loss of genetic variability (as measured by *N*_*e*_ and *U*_*W*_ ) reduced the total genetic variability (even though %GCA remained highest of all structures), while at the same time the other structures had enough time to build up enough genetic constraints to reach a %GCA sufficient for effective selection response. Thus, during the last third of the simulation, the centralized structure was superior for lower values of *K* and the distributed structure for intermediate to high complexity levels.

The pattern observed in the final cycles reflects our findings in our previous study (Technow *et al*. 2021), except that the relative performance of the structures in regards to *K* was shifted to higher values. For example, while in Technow *et al*. (2021) we found the centralized structure to be superior only until *K* = 4, we now found it to be superior until *K* = 7. Similarly, for the distributed structure which there was superior between *K* = 5–8 and here from about 8–13. This is of course because of the added dynamics of a shifting TPE, which increase the importance of preserving genetic variability for long-term genetic gain in order to react to the environmental shifts. The isolated structures struggled with this, and for lower to immediate values of K, where genetic variability had been largely exhausted by the final cycles (Figure 9), even experienced a decrease in overall performance level (Figure 5) as previously advantageous allele and gene complexes, that had become fixed, ultimately became neutral or detrimental. There is thus a balance between short and long-term selection response, which translates to the compromise between exposing and exploiting additive genetic variation within the current TPE and retaining genetic variability for adapting to the future TPE. In other words, GxE and a shifting TPE expand the complexity of the potential state space (i.e., the potentially relevant and accessible genotype configurations) and creates greater uncertainty associated with pursuing any specific selection trajectory. This confers an advantage for more landscape exploration before committing to move towards a particular local optimum. The different breeding program structures adjust the balance of exploration and exploitation differently. The distributed structure, which maintains separate sub-programs but with constant exchange between them, strikes a middle ground between the centralized and isolated structures and combined the desired features of both while mitigating some of their drawbacks. For example, it was able to quickly generate enough additive variation, measured as %GCA, for facilitating a selection response (Figure 7), but without the dramatic drop in genetic variability (Figure 9) and effective population size (Figure 8) experienced by the isolated structures. Across a wide range of degrees of complexity, this allowed it to quickly generate cycle over cycle performance improvements during the early cycles, while maintaining these responses also in the later cycles. As outlined in Technow *et al*. (2021), this distributed structure closely reflects the breeding program model that emerged organically over the last century within large scale hybrid breeding operations (Crabb 1947; Cooper *et al*. 2014).

**Figure 9.**
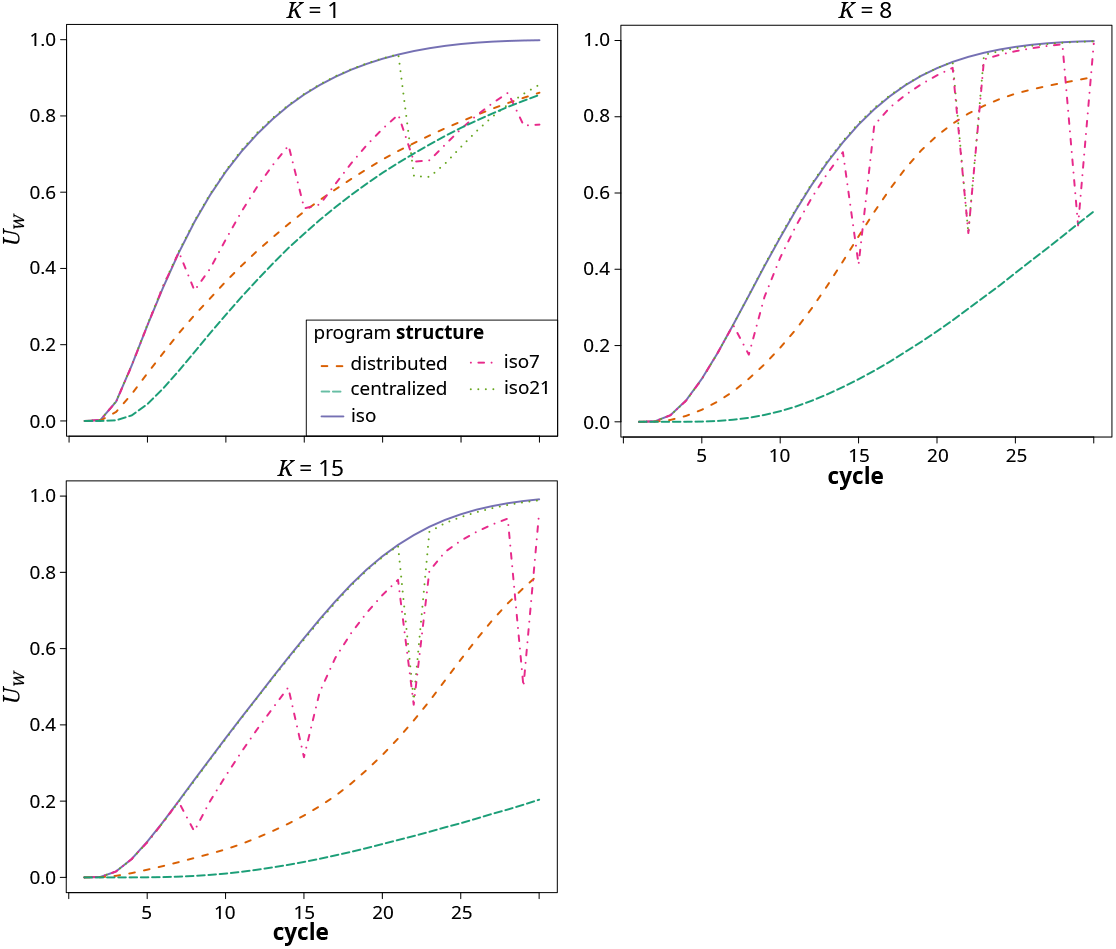
Proportion of loci with MAF *<* 0.05 (U-shapedness, *U*_*W*_ ) across cycles for the different breeding program structures in the gradual TPE change scenario at *K* values of 1 (top left), 8 (top right) and 15 (bottom left). The values represented are the averages across the replications of the simulation.

**Figure 10.**
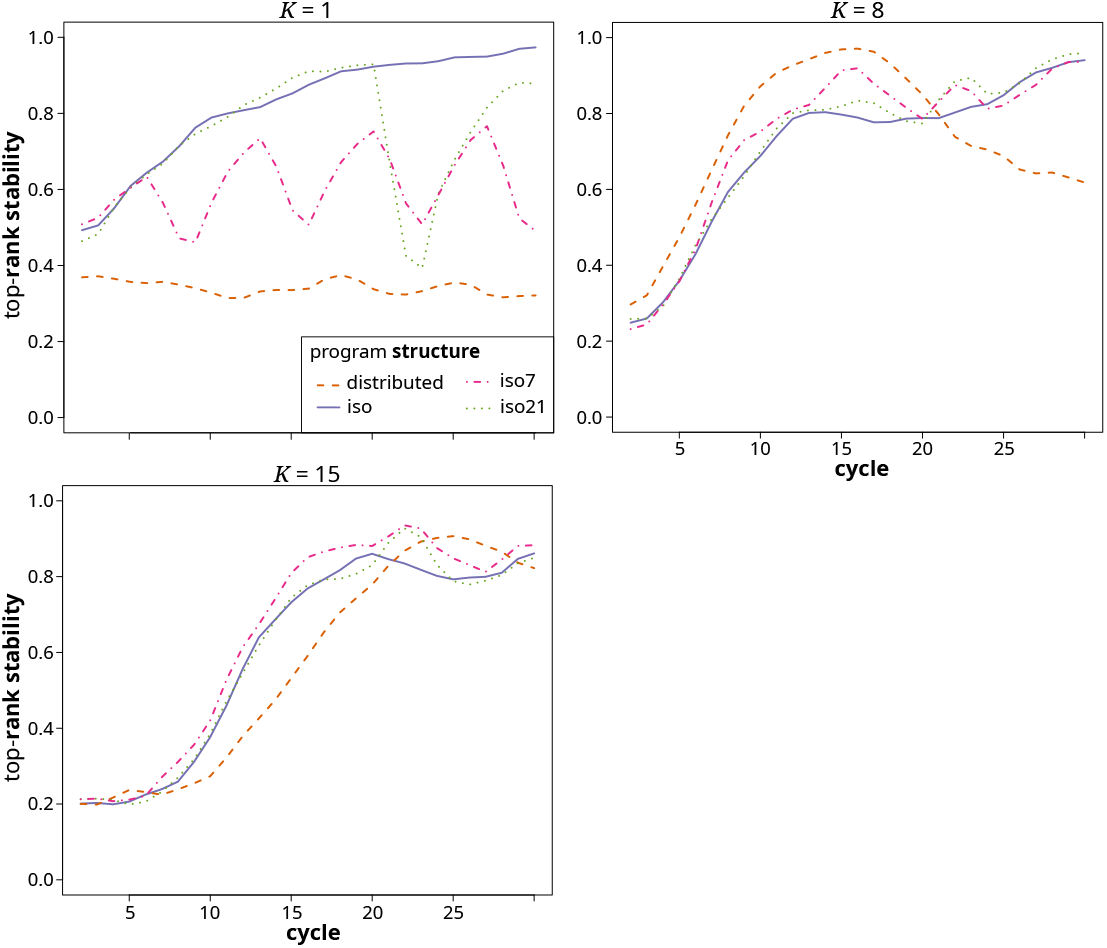
Top-rank stability (probability that the top-ranked program in cycle *n* was the same as the one in cycle *n* −1) across cycles for the different breeding program structures in the gradual TPE change scenario at *K* values of 1 (top left), 8 (top right) and 15 (bottom left). Note that this metric is not applicable to the centralized structure.

The notable exception to these trends is the special case of *K* = 1, where all variation is inherently additive and the outlined effects of population sub-division on the emergence of additive variation do not apply. Here the centralized structure, with its substantially higher *N*_*e*_, was superior throughout, as would be expected from well-established results pertaining to the assumptions of the infinitesimal model with additive effects (Barton *et al*. 2017).

### Restoring genetic variability through germplasm exchange

Restoring the genetic variability of isolated breeding programs in the public sector through occasional germplasm exchange was proposed as one option for ensuring their long-term adaptability and competitiveness (Fonseca *et al*. 2021). We evaluated this option in the form of the iso7 and iso21 structures that allowed for sporadic germplasm exchange between the otherwise isolated programs. Interestingly, these followed the performance development of the iso structure, in which no exchange takes place, closely. The germplasm exchange thus had little to no effect on the medium-to long-term behavior of the germplasm evolutionary trajectory. As can be seen for *U*_*W*_ (Figure 9), and, albeit less clearly, for *N*_*e*_ (Figure 8), the sporadic germplasm exchange did have a short term effect on genetic variability, which however dissipated very rapidly in a matter of a cycle or two. The explanation can be found in the near zero across program correlation of GCA effects in the isolated structures (Figure 7), which means that germplasm that is elite in one sub-group is not likely to be also elite in the other and hence will quickly be purged. As we pointed out here and previously (Technow *et al*. 2021), not just the (relative) amount of additive variation, but also the directionality and magnitude of additive effects are dependent on the overall germplasm context. Thus, the additive effect of a gene or gene complex in one context is not necessarily predictive of its (additive) effect in another. With the distributed structure, this GCA correlation started increasing as soon as a sizable amount of GCA variation was exposed. Owing to the constant exchange of germplasm, the sub-programs thus were converging towards a section of the genetic landscape within which additive genetic effects have at least some transferability from one genetic background to the next. Also of note is that for the distributed structure, the rank-stability metric in the final third of the breeding simulation was around 0.6, i.e., low enough to indicate frequent reranking of sub-programs. During the final phase of the simulation, where access to sources of genetic variability was most crucial, the direction of the flow of germplasm hence changed relatively frequently. Interestingly, both the maintenance of transferability of gene effects, as well as the fluidity of the ranking of the programs, existed despite considerable sub-population differentiation evidenced by the Fst statistic (Figure 8). The different sub-populations thus did not all occupy the same genetic space either, meaning that this structure indeed exhibited the features of a distributed search strategy in complex genetic space (Podlich and Cooper 1999). The relatively high Fst values are also indicative of the presence of significant amounts of latent genetic variability at the meta-population level, i.e., across the ensemble of sub-programs comprising the breeding structure (Wade and Goodnight 1998). This is even more the case for the isolated structures in which the very high Fst values indicated fixation or near fixation of contrasting gene complexes across the different sub-populations. However, in contrast to the distributed structure, the described difficulties with germplasm exchange meant that this latent genetic variability could not be utilized effectively.

The problem of incorporating unadapted and inferior germplasm for maintaining genetic variability has long been recognized and several strategies to mitigate undesirable effects on the recipient germplasm exist (Allier *et al*. 2020). Here we highlight that this problem does not just apply to exotic or ancient genetic resources derived from, e.g., gene banks (Yu *et al*. 2016) or landraces (Böhm *et al*. 2017), but also to sources that can be considered “elite” within their own genetic context and adapted to the same TPE. Usually it is recommended to adapt these genetic resources, elite or not, before their introgression with the elite germplasm pool using “pre-breeding”, which in the era of genomic selection can be done rapidly and cost effectively (Polzer *et al*. 2025). Because of the context dependency of genetic effects, it is important that this adaptation is done within the genetic context of the recipient germplasm (Fonseca *et al*. 2021).

In our simulation, the maximum amount of breeding crosses with external germplasm during the exchanges was 25%. The persistence and impact of the exchanged germplasm would have been greater at higher exchange rates, because in that case the germplasm context of the recipient program after the first recombination step would have been more similar to that of the donor program. In the extreme case, one could envision a strategy where the germplasm of the inferior programs is essentially replaced with that of the superior programs. This, however, is not common practice, as the resulting homogenization and loss of overall genetic variability would largely defeat the purpose of maintaining separate breeding programs. The rapid genetic variability loss in the isolated structures could potentially also have been mitigated by employing variability preserving selection techniques, such as optimal mate selection (Cowling *et al*. 2017) or techniques aiming to preserve haplotype diversity (Müller *et al*. 2018). Furthermore, we did not allow for the possibility of de novo generation of genetic variability, through e.g., mutations (Durand *et al*. 2010) or epigenetic phenomena (Hauben *et al*. 2009), which when propagated and magnified through complex gene interaction networks (Rasmusson and Phillips 1997), might facilitate the maintenance of selection response across many generations even in isolated and relatively narrow breeding populations (Dudley and Lambert 2010; Durand *et al*. 2010, 2015).

### Adaptation to environmental complexity and change

We considered two contrasting environmental change scenarios: gradual change, where the TPE changes slowly from one end of the environmental spectrum to the next, and sudden change, where a dramatic shift happens from one cycle to the next. Overall, we found that the trends in relative superiority of the different structures across cycles and complexity levels as well as for most metrics used to describe these trends, were very similar in principle across both scenarios. We also briefly considered a cyclical change scenario, where the TPE oscillates across the environmental spectrum and also here found the same general trends (results not shown). One explanation for this is that over the long-term, the TPE in both change scenarios traverses a similarly wide range of environment types, while in the short term there is little practical difference between the directional change in TPE weights in the gradual scenario and the small random fluctuations of TPE weights in the sudden scenario. This basic similarity is also apparent from the similar curves of lagged genotypic correlation between cycles (Figure 1).

A sudden shift was expected to represent a shock to the system that can create bigger short-term disturbances, like throwing a large rock into water, than the effects of the gradual change, which is more like changing the tide. In our simulation, however, this shift happened only after cycle 20, after which the genetic space had already linearized to a large degree. The emerged additivity thus apparently persisted regardless, which can be seen in that there was no noticeable drop in %GCA at that point (Figure 7). The structures thus quickly regained their selection trajectories and the dynamics prior to the shift continued. In other words, while the emerged additive gene effects might have changed in their relative magnitude and sign, the system remained linearized, as its baseline topology (the *N* and *K* parameters) and more importantly the effective topology, i.e., the number of still segregating loci within each interactive pathway, were not affected. However, environmental shifts can directly alter genetic and phenotypic expression through increases in mutation rate (Carja *et al*. 2014), induction of epigenetic changes (Turner 2009) or even modify the topology of biological networks through recruitment of dormant pathways or increases in their interconnectivity (Iwasaki *et al*. 2013). While not considered here, such phenomena could be readily incorporated into the *NK* framework (Altenberg 1994).

In the absence of sources of de novo genetic variation, it seems that what matters most for the short and long-term behavior of breeding germplasm, is whether or not the TPE changes, rather than how. As detailed above, the salient consequence of a shifting TPE is the increased importance of conserving genetic variability to facilitate adapting to future environmental conditions. In a static TPE and resulting static genetic landscape, a local optimum identified early persists. There is thus less penalty for quickly and severely constraining genetic space as under a shifting TPE and landscape, where a local optimum can disappear over time. Thus, whereas under a static TPE, an exhaustion of genetic variability will lead only to a slowing rate of performance improvements, under a dynamic TPE it can mean a decline in absolute performance level, as observed for the isolated structures under low to intermediate levels of complexity (Figure 5). Regardless of whether the long-term environmental shift occurs gradually or suddenly, environmental conditions continuously change and fluctuate across all time scales (Bernhardt *et al*. 2020). This was represented in our simulations by small random cycle over cycle variations in TPE weights. These short-term fluctuations might in fact help to conserve genetic variability (Bell 2010; Abdul-Rahman *et al*. 2021), because the small and non-directional changes in the underlying genetic landscape continuously perturb selection trajectories and thus prevent or slow down the premature fixation of gene complexes that might become disadvantageous later.

The threat of climate change and the renewed appreciation for the role of GxE have led to an increased awareness in the plant breeding community for the need to enable germplasm to adapt to a the changing and likely more challenging environmental and more constrained agronomic management options of the future. This has also led to an awareness of the need to align the multi-environment field trials (MET) in which the germplasm is evaluated and selected to conditions it ultimately will have to perform under in the on-farm TPE (Cooper *et al*. 2022). This has not always been consensus, as plant breeders historically often preferred the increased precision and convenience of a network of well-managed, homogeneous and high yielding locations within a fixed research network, over trialing under the more realistic but also more challenging conditions encountered on-farm, particularly concerning environmental stress and input limitations (Ceccarelli and Grando 1996; Bänziger and Cooper 2001). Our simulation assumed that the MET captured the different environment types in the exact frequency with which they occur in the TPE. This is not unrealistic, considering novel multi-environment trial designs facilitated by whole genome prediction, such as sparse location testing (Jarquin *et al*. 2020), possibly even directly on participatory farms (Werner *et al*. 2025), a deepened appreciation for the need of constructing appropriate METs (Cooper *et al*. 2022), and methods for removing remaining distributional biases through appropriate reweighing of environment type frequencies in the MET (Podlich *et al*. 1999; Chenu 2015).

Beyond simply following the trajectory of the TPE, adaptation to future environmental conditions or management regimes can be anticipated by representing these in the MET, either by up-weighing the corresponding environment types, evaluating and selecting the germplasm in managed stress environments (Bänziger and Cooper 2001) or virtually through predictive models (Technow *et al*. 2015; Messina *et al*. 2018). An even more explicit approach is selecting for designed ideotypes with physiological features expected to convey adaptation to the anticipated conditions (Semenov and Stratonovitch 2013; Hammer *et al*. 2020). Evaluating today’s germplasm under hypothetical future conditions can be valuable for identifying weaknesses that could critically limit adaptability to these conditions. However, as these efforts require resources, care should be taken that the *adjacent possible* (Kauffman 2014) performance under current and near-term conditions does not become an afterthought. Not least because of the inherent uncertainty involved in anticipating environmental conditions decades into the future.

We already highlighted the distributed structure as achieving the best balance between short and long-term genetic gain. Under a changing TPE, with uncertain long-term future conditions, this property can also help to ensure adaptation to a wide array of possibilities, simply by not just distributing the germplasm under evaluation but also the environments under which it is evaluated. This in fact has been common practice in breeding programs in large organizations, where rarely two programs target entirely overlapping environmental zones. The diverging selection environments will reinforce the implications of a stratified germplasm space under complex trait genetics. For the isolated structures it will likely mean an even faster genetic divergence, which in combination with the different selection environment will make the highlighted issues with germplasm exchange even more challenging. On the other hand, for a distributed structure with regular exchange, it could have the positive effect of retaining more (latent) genetic variability on the meta-population level as well as exploration of more unique physiological adaptation strategies. An interesting novel concept combining a distributed germplasm strategy with environmental stratification through participatory on-farm field experimentation was recently proposed by Santamarina *et al*. (2025) for the purpose of developing locally adapted crop mixtures for complex TPE.

### A framework for studying the short and long-term behavior of breeding programs

As in our previous work, our objective was not to provide specific and ultimate answers as to the optimal design of breeding programs but to suggest a quantitative framework, comprising of a model for biological complexity and breeding as well as metrics for describing their behavior, under which concrete questions can be studied. This framework is based on the *E(NK)* model (Cooper and Podlich 2002), which is an extension of the *NK* model. The latter was developed by the theoretical biologist and complexity researcher Stuart A. Kauffman (Kauffman 1993) to study evolution in complex biological systems. *NK*, respectively *E(NK)*, models are an attractive choice because they allow to generate genetic landscapes with finely tunable degrees of complexity, from the special case of *K* = 1 representing intrinsic and complete additivity to almost intractable complexity at values of *K* = 15 and beyond (Figure 3). Another advantage is that *NK* models are relatively efficient to represent and evaluate computationally, even at genome scale levels of dimensionality and interconnectedness, when using techniques such as random functions (Altenberg 1994). Finally, the *NK* model has established itself as a representative model of complex systems beyond biology. It can thus provide a canonical framework across a wide array of disciplines concerned with search strategies in complex systems, including consumer behavior research (Li *et al*. 2022), innovation management (Ganco 2017), physics (Qu *et al*. 2002), infrastructure design (Grove and Baumann 2012) and economics (Faggini and Parziale 2016).

A limitation of the generic nature of the *E(NK)* framework is that it allows only to address conceptual “what if” questions. For practical plant breeding operations *E(NK)* like models that reflect the features and topology of their actual germplasm space and TPE would be of great interest. For model organisms with simple genomes and in clearly defined environments, it is theoretically possible to directly estimate topological features of *NK* like fitness landscapes through studying the effects of individual mutations on phenotypes and from those make predictions of, e.g., accessible evolutionary trajectories (de Visser and Krug 2014). However, for agricultural crop species, with complex genomes and exposure to TPE landscape characterized by complex GxE interactions, such a direct approach seems infeasible. A more practical approach is to use biological models, which instead of attempting to directly link genome features to phenotypic variation, represent, on the basis of prior biological experimentation, simpler and intermediate hier-archical levels in the gene to phenotype map (Hammer *et al*. 2006, 2019). A prime example in an agricultural context are dynamic crop growth models. They represent the biological knowledge gained from decades of research on the interplay between plant physiology, soil science and micro-meteorology (e.g., Keating *et al*. 2003; van Ittersum *et al*. 2003). With the help of Bayesian hierarchical whole genome regression techniques (Technow *et al*. 2015; Messina *et al*. 2018; Powell *et al*. 2021), these models have successfully been applied to model germplasm-specific biological complexity at the scale and resolution required by practical plant breeding operations (e.g., Cooper *et al*. 2016; Onogi 2020; Diepenbrock *et al*. 2022; Jighly *et al*. 2023). This approach has also been called “BioWGP”. On the opposite end of the gene to phenotype map from crop growth models exist gene regulatory networks models, which also have been applied in an agricultural and plant breeding context (Dong *et al*. 2012; Leong *et al*. 2025). They represent the complex relationships between genes, transcription factors and environmental signals that regulate gene expression and are constructed from a combination of experimentally obtained ‘omics’ data and prior information (Muhammad *et al*. 2017; Badia-i Mompel *et al*. 2023). The structure and properties of gene regulatory networks have been conceptualized in the *omnigenic* model (Boyle *et al*. 2017), which shares many features with the *NK* modeling framework and postulates that the polygenic nature of most quantitative traits is a consequence of effect propagation through these highly interconnected networks. Recently, Ružičková *et al*. (2024) developed a statistical approach that allows to apply this omnigenic model directly to empirical data sets. Their *quantitative omnigenic model* could be used to obtain germplasm specific estimates of allele effects within the general interactive network structure, similarly as the BioWGP approach, and thereby generate an *E(NK)*-like model of biological complexity tailored to a specific germplasm context. Crop growth models, gene regulatory networks and other biological models, such as metabolic networks (Fiévet *et al*. 2010), used individually or combined in dynamic gene to phenotype systems (Kauffman 2004; Messina *et al*. 2025), could thus be deployed to reduce to practice the conceptual insights gained from the *E(NK)* framework. If such models can capture the system topology, level of complexity and emergent behavior underlying an existing breeding germplasm, optimizing long-term breeding strategies for specific germplasm and TPE scenarios would be conceivable.

### Simple solutions to very complex problems?

The most basic result from our work is that the optimal breeding strategy for balancing short and long-term genetic gain depends almost entirely on the level of biological complexity inherent in the system. In particular we want to emphasize that ours and others results show that germplasm trajectories and behavior for the special case of pure additivity (*K* = 1) can be dramatically different from even low to intermediate levels of non-additivity (Rasmusson and Phillips 1997; Technow *et al*. 2021; Tessele *et al*. 2025). Biological complexity is an undeniable reality, even if it doesn’t always express itself in the form of statistically detectable non-additive variation (Huang and Mackay 2016). Current attention to optimizing genetic gain and managing genetic variation for the long term based implicitly on *K* = 1 additive and stationary infinitesimal type models within an often static TPE, does not adequately consider the details of GxE selection trajectories the breeding program has to navigate under a combination of genetic complexity and a shifting environment.

In an opinion paper, renowned plant breeding scientist Major M. Goodman (Goodman 2002) linked the recurrence of so called plant breeding “bandwagons” (Simmonds 1991; Bernardo 2016), i.e., technologies and approaches that often promise simple solutions for complex problems, to an under appreciation of GxE interaction and epistasis (i.e., biological complexity and context dependency of genetic effects). We believe that novel technologies can play an important role in helping plant breeders tackle the challenges of future climates and population demands. However, their effective use requires a framework to appropriately represent the biological complexity and emergent phenomena of the genetic and environmental space in which they hope to intervene. Such a framework must also be able to represent the self-organizing principles with which germplasm has successfully navigated challenging and complex GxE landscapes over the last century, and if structured and managed appropriately, can do so also in the future.

## Literature cited

Abdul-Rahman, F., D. Tranchina, and D. Gresham, 2021 Fluctuating environments maintain genetic diversity through neutral fitness effects and balancing selection. Mol Biol Evol 38: 4362–4375.

Allier, A., S. Teyssèdre, C. Lehermeier, L. Moreau, and A. Charcosset, 2020 Optimized breeding strategies to harness genetic resources with different performance levels. BMC Genomics 21: 349.

Altenberg, L., 1994 Evolving better representations through selective genome growth. In Proceedings of the First Ieee Conference on Evolutionary Computation IEEE World Congress on Computational Intelligence ICEC-94, pp. 182–187, IEEE.

Atlin, G. N., J. E. Cairns, and B. Das, 2017 Rapid breeding and varietal replacement are critical to adaptation of cropping systems in the developing world to climate change. Glob Food Sec 12: 31–37.

Badia-i Mompel, P., L. Wessels, S. Müller-Dott, R. Trimbour, R. O. Ramirez Flores, et al., 2023 Gene regulatory network inference in the era of single-cell multi-omics. Nat Rev Genet 24: 739–754.

Baker, L. H. and R. N. Curnow, 1969 Choice of Population Size and Use of Variation Between Replicate Populations in Plant Breeding Selection Programs 1. Crop Sci 9: 555–560.

Bänziger, M. and M. Cooper, 2001 Breeding for low input conditions and consequences for participatory plant breeding examples from tropical maize and wheat. Euphytica 122: 503–519.

Barton, N. H., A. M. Etheridge, and A. Véber, 2017 The infinitesimal model: Definition, derivation, and implications. Theor Pop Biol 118: 50–73.

Basso, B., A. D. Kendall, and D. W. Hyndman, 2013 The future of agriculture over the Ogallala Aquifer: Solutions to grow crops more efficiently with limited water. Earths Future 1: 39–41.

Becklin, K. M., J. T. Anderson, L. M. Gerhart, S. M. Wadgymar, C. A. Wessinger, et al., 2016 Examining plant physiological responses to climate change through an evolutionary lens. Plant Physiol 172: 635–649.

Bell, G., 2010 Fluctuating selection: the perpetual renewal of adaptation in variable environments. Philos Trans R Soc Lond B Biol Sci 365: 87–97.

Bernardo, R., 2016 Bandwagons I, too, have known. Theor Appl Genet 129: 2323–2332.

Bernhardt, J. R., M. I. O’Connor, J. M. Sunday, and A. Gonzalez, 2020 Life in fluctuating environments. Philos Trans R Soc B 375: 20190454.

Böhm, J., W. Schipprack, H. F. Utz, and A. E. Melchinger, 2017 Tapping the genetic diversity of landraces in allogamous crops with doubled haploid lines: a case study from European flint maize. Theor Appl Genet 130: 861–873.

Boichard, D., L. Maignel, and É. Verrier, 1997 The value of using probabilities of gene origin to measure genetic variability in a population. Genet Sel Evol 29: 5–23.

Boyle, E. A., Y. I. Li, and J. K. Pritchard, 2017 An expanded view of complex traits: from polygenic to omnigenic. Cell 169: 1177–1186.

Brown, J. K. M. and J. C. Rant, 2013 Fitness costs and trade-offs of disease resistance and their consequences for breeding arable crops. Plant Pathol 62: 83–95.

Burdon, J. J. and P. H. Thrall, 2003 The fitness costs to plants of resistance to pathogens. Genome Biol 4: 227.

Carja, O., U. Liberman, and M. W. Feldman, 2014 Evolution in changing environments: Modifiers of mutation, recombination, and migration. Proc Nat Acad Sci 111: 17935–17940.

Carlborg, O. and C. S. Haley, 2004 Epistasis: too often neglected in complex trait studies? Nat Rev Genet 5: 618–625.

Ceccarelli, S. and S. Grando, 1996 Drought as a challenge for the plant breeder. Plant growth regulation 20: 149–155.

Chapman, S. C., S. Chakraborty, M. F. Dreccer, and S. M. Howden, 2012 Plant adaptation to climate change—opportunities and priorities in breeding. Crop Pasture Sci 63: 251–268.

Chenu, K., 2015 Characterizing the crop environment – nature, significance and applications. In Crop Physiology (Second Edition), pp. 321–348, Academic Press, San Diego.

Clements, H. F., 1964 Interaction of factors affecting yield. Annu Rev Plant Phys 15: 409–442.

Collins, B. and K. Chenu, 2021 Improving productivity of Australian wheat by adapting sowing date and genotype phenology to future climate. Clim Risk Manag 32: 100300.

Comstock, R. and R. H. Moll, 1963 Genotype-environment interactions. Statistical genetics and plant breeding 982: 164–196.

Comstock, R. E., 1977 Quantitative genetics and the design of breeding programs. In Proceedings of the International Conference on Quantitative Genetics, pp. 16–21, Ames, IA, Iowa State University Press.

Cooper, M. and I. H. DeLacy, 1994 Relationships among analytical methods used to study genotypic variation and genotype-by-environment interaction in plant breeding multi-environment experiments. Theor Appl Genet 88: 561–572.

Cooper, M. and C. D. Messina, 2023 Breeding crops for drought-affected environments and improved climate resilience. The Plant Cell 35: 162–186.

Cooper, M., C. D. Messina, D. Podlich, L. R. Totir, A. Baumgarten, et al., 2014 Predicting the future of plant breeding: complementing empirical evaluation with genetic prediction. Crop Pasture Sci 65: 311.

Cooper, M., C. D. Messina, T. Tang, C. Gho, O. M. Powell, et al., 2022 Predicting Genotype × Environment × Management (G × E × M) interactions for the design of crop improvement strategies: integrating breeder, agronomist, and farmer perspectives. In Plant Breeding Reviews, edited by I. Goldman, pp. 467–585, Wiley.

Cooper, M. and D. W. Podlich, 2002 The E(NK) model: extending the NK model to incorporate gene-by-environment interactions and epistasis for diploid genomes. Complexity 7: 31–47.

Cooper, M., D. W. Podlich, and O. S. Smith, 2005 Gene-to-phenotype models and complex trait genetics. Aust J Agric Res 56: 895–918.

Cooper, M., F. Technow, C. Messina, C. Gho, and L. R. Totir, 2016 Use of crop growth models with whole-genome prediction: application to a maize multienvironment trial. Crop Sci 56: 2141–2156.

Cooper, M., K. P. Voss-Fels, C. D. Messina, T. Tang, and G. L. Hammer, 2021 Tackling G × E × M interactions to close on-farm yield-gaps: creating novel pathways for crop improvement by predicting contributions of genetics and management to crop productivity. Theor Appl Genet 134: 1625–1644.

Corbin, L. J., A. Y. H. Liu, S. C. Bishop, and J. A. Woolliams, 2012 Estimation of historical effective population size using linkage disequilibria with marker data. J Anim Breed Genet 129: 257–270.

Cowling, W. A., L. Li, K. H. M. Siddique, M. Henryon, P. Berg, et al., 2017 Evolving gene banks: improving diverse populations of crop and exotic germplasm with optimal contribution selection. J Exp Bot 68: 1927–1939.

Crabb, A. R., 1947 The Hybrid-Corn Makers: Prophets of Plenty. Rutkes University Press.

Csaszar, F. A., 2018 A note on how NK landscapes work. J Org Design 7: 10.1186/s41469–018–0039–0.

de Visser, J. A. G. M. and J. Krug, 2014 Empirical fitness landscapes and the predictability of evolution. Nat Rev Genet 15: 480–490.

Diepenbrock, C. H., T. Tang, M. Jines, F. Technow, S. Lira, et al., 2022 Can we harness digital technologies and physiology to hasten genetic gain in US maize breeding? Plant Physiol 188: 1141–1157.

Dong, Z., O. Danilevskaya, T. Abadie, C. Messina, N. Coles, et al., 2012 A gene regulatory network model for floral transition of the shoot apex in maize and its dynamic modeling. PLoS ONE 7: e43450.

Dudley, J. W. and R. J. Lambert, 2010 100 Generations of Selection for Oil and Protein in Corn. In Plant Breeding Reviews, pp. 79–110, John Wiley & Sons, Ltd.

Durand, E., M. I. Tenaillon, X. Raffoux, S. Thépot, M. Falque, et al., 2015 Dearth of polymorphism associated with a sustained response to selection for flowering time in maize. BMC Evol Biol 15: 10.1186/s12862–015–0382–5.

Durand, E., M. I. Tenaillon, C. Ridel, D. Coubriche, P. Jamin, et al., 2010 Standing variation and new mutations both contribute to a fast response to selection for flowering time in maize inbreds. BMC Evol Biol 10: 10.1186/1471–2148–10–2.

Duvick, D., 1999 Heterosis: feeding people and protecting natural resources. In The genetics and exploitation of heterosis in crops, edited by J. Coors and S. Pandey, pp. 19–29, CSSA, Madison, WI.

Duvick, D., J. Smith, and M. Cooper, 2004 Long-term selection in a commercial hybrid maize breeding program. In Plant Breeding Reviews, edited by J. Janick, pp. 109–152, John Wiley & Sons, Inc., Hoboken, NJ.

Dwivedi, S. L., A. B. Britt, L. Tripathi, S. Sharma, H. D. Upadhyaya, et al., 2015 Haploids: constraints and opportunities in plant breeding. Biotechnol Adv 33: 812–829.

Elias, A. A., K. R. Robbins, R. Doerge, and M. R. Tuinstra, 2016 Half a century of studying Genotype × Environment interactions in plant breeding experiments. Crop Sci 56: 2090–2105.

Faggini, M. and A. Parziale, 2016 A new perspective for fiscal federalism: the NK model. J Econ Issues 50: 1069–1104.

Falconer, D. S., 1952 The problem of environment and selection. Am Nat 86: 293–298.

Fiévet, J. B., C. Dillmann, and D. de Vienne, 2010 Systemic properties of metabolic networks lead to an epistasis-based model for heterosis. Theor Appl Genet 120: 463–473.

Fiévet, J. B., T. Nidelet, C. Dillmann, and D. de Vienne, 2018 Heterosis is a systemic property emerging from non-linear genotype-phenotype relationships: evidence from in vitro genetics and computer simulations. Front Genet 9: 10.3389/fgene.2018.00159.

Fonseca, J. M. O., R. Perumal, P. E. Klein, R. R. Klein, and W. L. Rooney, 2021 Combining abilities and elite germplasm enhancement across U.S. public sorghum breeding programs. Crop Sci 61: 4098–4111.

Galluzzi, G., A. Seyoum, M. Halewood, I. López Noriega, and E. W. Welch, 2020 The role of genetic resources in breeding for climate change: the case of public breeding programmes in eighteen developing countries. Plants 9: 1129.

Ganco, M., 2017 NK model as a representation of innovative search. Research Policy 46: 1783–1800.

Gao, L., M. B. Kantar, D. Moxley, D. Ortiz-Barrientos, and L. H. Rieseberg, 2023 Crop adaptation to climate change: an evolutionary perspective. Molecular Plant 16: 1518–1546.

Gauch Jr., H. G. and R. W. Zobel, 1997 Identifying mega-environments and targeting genotypes. Crop Sci 37: 311–326.

Goodman, M. M., 2002 New sources of germplasm: Lines, transgenes, and breeders. In Mem. Congresso Nacional de Fitogenetica, pp. 28–41, Univ. Autonimo Agr. Antonio Narro, Saltillo, Coah. Mexico.

Goudet, J. and T. Jombart, 2022 hierfstat: estimation and tests of hierarchical F-Statistics. R package version 0. 5-11.

Grove, N. and O. Baumann, 2012 Complexity in the telecommunications industry: When integrating infrastructure and services backfires. Telecomm Policy 36: 40–50.

Guo, M., M. A. Rupe, J. Wei, C. Winkler, M. Goncalves-Butruille, et al., 2014 Maize ARGOS1 (ZAR1) transgenic alleles increase hybrid maize yield. J Exp Bot 65: 249–260.

Hadfield, J. D., 2010 Mcmc methods for multi-response generalized linear mixed models: The MCMCglmm R package. J Stat Soft 33: 1–22.

Hammer, G., M. Cooper, F. Tardieu, S. Welch, B. Walsh, et al., 2006 Models for navigating biological complexity in breeding improved crop plants. Trends Plant Sci 11: 587–593.

Hammer, G., C. Messina, A. Wu, and M. Cooper, 2019 Biological reality and parsimony in crop models—why we need both in crop improvement! in silico Plant 1: diz010.

Hammer, G. L., G. McLean, E. van Oosterom, S. Chapman, B. Zheng, et al., 2020 Designing crops for adaptation to the drought and high-temperature risks anticipated in future climates. Crop Sci 60: 605–621.

Harrington, S. A., A. E. Backhaus, A. Singh, K. Hassani-Pak, and C. Uauy, 2020 The wheat GENIE3 network provides biologically-relevant information in polyploid wheat. G3 10: 3675–3686.

Hauben, M., B. Haesendonckx, E. Standaert, K. Van Der Kelen, A. Azmi, et al., 2009 Energy use efficiency is characterized by an epigenetic component that can be directed through artificial selection to increase yield. Proc Natl Acad Sci 106: 20109–20114.

Heffner, E. L., A. J. Lorenz, J.-L. Jannink, and M. E. Sorrells, 2010 Plant breeding with genomic selection: gain per unit time and cost. Crop sci 50: 1681–1690.

Hill, W. G., M. E. Goddard, and P. M. Visscher, 2008 Data and theory point to mainly additive genetic variance for complex traits. PLoS Genet 4: e1000008.

Huang, W. and T. F. C. Mackay, 2016 The genetic architecture of quantitative traits cannot be inferred from variance component analysis. PLOS Genet 12: 10.1371/journal.pgen.1006421.

Iwasaki, W. M., M. E. Tsuda, and M. Kawata, 2013 Genetic and environmental factors affecting cryptic variations in gene regulatory networks. BMC Evol Biol 13: 91.

Jarquin, D., R. Howard, J. Crossa, Y. Beyene, M. Gowda, et al., 2020 Genomic prediction enhanced sparse testing for multi-environment trials. G3 10: 2725–2739.

Jighly, A., T. Thayalakumaran, G. J. O’Leary, S. Kant, J. Panozzo, et al., 2023 Using genomic prediction with crop growth models enables the prediction of associated traits in wheat. J Exp Bot 74: 1389–1402.

Kanter, D. R., O. Chodos, O. Nordland, M. Rutigliano, and W. Winiwarter, 2020 Gaps and opportunities in nitrogen pollution policies around the world. Nat Sustain 3: 956–963.

Karavolias, N. G., W. Horner, M. N. Abugu, and S. N. Evanega, 2021 Application of gene editing for climate change in agriculture. Front Sustain Food Syst 5: 685801.

Kauffman, S., 2004 A proposal for using the ensemble approach to understand genetic regulatory networks. J Theor Biol 230: 581–590.

Kauffman, S. A., 1993 The Origins of Order: Self-Organization and Selection in Evolution. Oxford University Press, New York.

Kauffman, S. A., 2014 Prolegomenon to patterns in evolution. Biosystems 123: 3–8.

Kauffman, S. A. and E. D. Weinberger, 1989 The NK model of rugged fitness landscapes and its application to maturation of the immune response. J Theor Biol 141: 211–245.

Keating, B. A., P. S. Carberry, G. L. Hammer, M. E. Probert, M. J. Robertson, et al., 2003 An overview of APSIM, a model designed for farming systems simulation. Eur J Agron 18: 267–288.

Kusmec, A., Z. Zheng, S. Archontoulis, B. Ganapathysubramanian, G. Hu, et al., 2021 Interdisciplinary strategies to enable data-driven plant breeding in a changing climate. One Earth 4: 372–383.

Leong, R., X. He, B. S. Beijen, T. Sakai, J. Goncalves, et al., 2025 Unlocking gene regulatory networks for crop resilience and sustainable agriculture. Nat Biotechnol in press: doi:10.1038/s41587–025–02727–4.

Li, X., T. Guo, Q. Mu, X. Li, and J. Yu, 2018 Genomic and environmental determinants and their interplay underlying phenotypic plasticity. Proc Natl Acad Sci 115: 6679–6684.

Li, X., T. Sekiguchi, J. Wu, and Q. Ye, 2022 Computational modeling of the value co-creation process in customer service: an application of the NK model. Front Psychol 13: 868803.

Löffler, C. M., J. Wei, T. Fast, J. Gogerty, S. Langton, et al., 2005 Classification of maize environments using crop simulation and geographic information systems. Crop Sci 45: 1708–1716.

Mayer, M., A. C. Hölker, E. González-Segovia, E. Bauer, T. Presterl, et al., 2020 Discovery of beneficial haplotypes for complex traits in maize landraces. Nat Commun 11: 4954.

McCann, H. C., 2020 Skirmish or war: the emergence of agricultural plant pathogens. Curr Opin Plant Biol 56: 147–152.

Melchinger, A. E., H. H. Geiger, G. Seitz, and G. A. Schmidt, 1987 Optimum prediction of three-way crosses from single crosses in forage maize (Zea mays L.). Theor Appl Genet 74: 339–345.

Melchinger, A. E. and R. K. Gumber, 1998 Overview of heterosis and heterotic groups in agronomic crops. In Concepts and Breeding of Heterosis in Crop Plant, pp. 29–44, CSSA.

Messina, C., J. Garcia-Abadillo, O. Powell, S. Tomura, A. Zare, et al., 2025 Toward a general framework for AI-enabled prediction in crop improvement. Theor Appl Genet 138: 151.

Messina, C., F. Technow, T. Tang, R. Totir, C. Gho, et al., 2018 Leveraging biological insight and environmental variation to improve phenotypic prediction: Integrating crop growth models (CGM) with whole genome prediction (WGP). Eur J Agron 100: 151–162.

Meuwissen, T. H. E., B. J. Hayes, and M. E. Goddard, 2001 Prediction of total genetic value using genome-wide dense marker maps. Genetics 157: 1819–1829.

Mikel, M. A. and J. W. Dudley, 2006 Evolution of North American dent corn from public to proprietary germplasm. Crop Sci 46: 1193–1205.

Milocco, L. and I. Salazar-Ciudad, 2022 Evolution of the G matrix under nonlinear genotype-phenotype maps. Am Nat 199: 420–435.

Montana, G., 2005 Hapsim: a simulation tool for generating haplotype data with pre-specified allele frequencies and ld coefficients. Bioinformatics 21: 4309–4311.

Moore, V. M., T. Peters, B. Schlautman, and E. C. Brummer, 2023 Toward plant breeding for multicrop systems. Proc Natl Acad Sci 120: e2205792119.

Muhammad, D., S. Schmittling, C. Williams, and T. A. Long, 2017 More than meets the eye: emergent properties of transcription factors networks in Arabidopsis. Biochim Biophys Acta 1860: 64–74.

Müller, D., P. Schopp, and A. E. Melchinger, 2018 Selection on expected maximum haploid breeding values can increase genetic gain in recurrent genomic selection. G3 8: 1173–1181.

Onogi, A., 2020 Connecting mathematical models to genomes: Joint estimation of model parameters and genome-wide marker effects on these parameters. Bioinformatics p. btaa129.

Peng, B., K. Guan, J. Tang, E. A. Ainsworth, S. Asseng, et al., 2020 Towards a multiscale crop modelling framework for climate change adaptation assessment. Nat Plants 6: 338–348.

Phillips, P. C., 2008 Epistasis – the essential role of gene interactions in the structure and evolution of genetic systems. Nat Rev Genet 9: 855–867.

Piepho, H. P., 1998 Methods for comparing the yield stability of cropping systems. J Agron Crop Sci 180: 193–213.

Podlich, D. and M. Cooper, 1999 Modelling plant breeding programs as search strategies on a complex response surface. Lecture Notes in Computer Science 1585: 171–178.

Podlich, D. W., M. Cooper, K. E. Basford, and H. H. Geiger, 1999 Computer simulation of a selection strategy to accommodate genotype environment interactions in a wheat recurrent selection programme. Plant Breed 118: 17–28.

Polzer, C., H.-J. Auinger, M. Terán-Pineda, A. C. Hölker, M. Mayer, et al., 2025 Rapid cycling genomic selection in maize landraces. Theor Appl Genet 138: 75.

Powell, O. M., F. Barbier, K. P. Voss-Fels, C. Beveridge, and M. Cooper, 2022 Investigations into the emergent properties of gene-to-phenotype networks across cycles of selection: a case study of shoot branching in plants. in silico Plant 4: diac006.

Powell, O. M., K. P. Voss-Fels, D. R. Jordan, G. Hammer, and M. Cooper, 2021 Perspectives on applications of hierarchical gene-to-phenotype (G2P) maps to capture non-stationary effects of alleles in genomic prediction. Front Plant Sci 12: 663565.

Press, W. H., S. A. Teukolsky, W. T. Vetterling, and B. P. Flannery, 1992 Numerical Recipes in C: The Art of Scientific Computing. Cambridge University Press, Cambridge.

Qu, X., M. Aldana, and L. P. Kadanoff, 2002 Numerical and theoretical studies of noise effects in the Kauffman model. J Stat Phys 109: 967–986.

R Core Team, 2021 R: a language and environment for statistical computing. R Foundation for Statistical Computing, Vienna, Austria.

Ramasubramanian, V. and W. D. Beavis, 2021 Strategies to assure optimal trade-offs among competing objectives for the genetic improvement of soybean. Front Genet 12: 675500.

Rasmusson, D. C. and R. L. Phillips, 1997 Plant breeding progress and genetic diversity from de novo variation and elevated epistasis. Crop Sci 37: 303–310.

Reif, J. C., A. E. Melchinger, and M. Frisch, 2005 Genetical and mathematical properties of similarity and dissimilarity coefficients applied in plant breeding and seed bank management. Crop Sci 45: 1–7.

Ružičková, N., M. Hledík, and G. Tkačik, 2024 Quantitative omnigenic model discovers interpretable genome-wide associations. Proc Nat Acad Sci 121: e2402340121.

Saha, R., P. F. Suthers, and C. D. Maranas, 2011 Zea mays iRS1563: a comprehensive genome-scale metabolic reconstruction of maize metabolism. PLoS ONE 6: 10.1371/journal.pone.0021784.

Santamarina, C., L. Mathieu, E. Bitocchi, A. Pieri, E. Bellucci, et al., 2025 Agroecological genomics and participatory science: optimizing crop mixtures for agricultural diversification. Trend Plant Sci 40382279: in press.

Schopp, P., C. Riedelsheimer, H. F. Utz, C.-C. Schön, and A. E. Melchinger, 2015 Forecasting the accuracy of genomic prediction with different selection targets in the training and prediction set as well as truncation selection. Theor Appl Genet 128: 2189–2201.

Semenov, M. A. and P. Stratonovitch, 2013 Designing high-yielding wheat ideotypes for a changing climate. Food and Energy Security 2: 185–196.

Seye, A. I., C. Bauland, A. Charcosset, and L. Moreau, 2020 Revisiting hybrid breeding designs using genomic predictions: simulations highlight the superiority of incomplete factorials between segregating families over topcross designs. Theor Appl Genet 133: 1995–2010.

Shi, Y., S. Pan, Y. You, S. A. Prior, D. Tian, et al., 2025 Extreme dry-heat climate impacts on greenhouse gas emission intensity in wheat production: insights and mitigation strategies. Glob Chang Biol 31: e70349.

Shull, G. H., 1908 The composition of a field of maize. J Hered os-4: 296–301.

Silva Dias, J. C., 2010 Impact of improved vegetable cultivars in overcoming food insecurity. Euphytica 176: 125–136.

Simmonds, N. W., 1991 Bandwagons I have known. TAA Newsletter pp. 7–11.

Simmons, C. R., H. R. Lafitte, K. S. Reimann, N. Brugière, K. Roesler, et al., 2021 Successes and insights of an industry biotech program to enhance maize agronomic traits. Plant Sci 307: 110899.

Sprague, G. F. and L. A. Tatum, 1942 General vs. specific combining ability in single crosses of corn. Agron J 34: 923–932.

Studer, A. J., H. Wang, and J. F. Doebley, 2017 Selection during maize domestication targeted a gene network controlling plant and inflorescence architecture. Genetics 207: 755–765.

Technow, F., 2013 hypred: simulation of genomic data in applied genetics. R package, version 0.4.

Technow, F. and J. P. Gerke, 2017 Parent-progeny imputation from pooled samples for cost-efficient genotyping in plant breeding. PLOS ONE 12: e0190271.

Technow, F., C. D. Messina, L. R. Totir, and M. Cooper, 2015 Integrating crop growth models with whole genome prediction through approximate bayesian computation. PLOS ONE 10: e0130855.

Technow, F., D. Podlich, and M. Cooper, 2021 Back to the future: implications of genetic complexity for the structure of hybrid breeding programs. G3 11: jkab153.

Tessele, A., G. Morris, E. Akhunov, B. Johnson, M. Clinesmith, et al., 2025 Interchromosomal linkage disequilibrium analysis reveals strong indications of sign epistasis in wheat breeding families. bioRxiv doi: 10.1101/2025.05.16.654524.

Tomura, S., O. Powell, M. J. Wilkinson, and M. Cooper, 2025 Ensemble-based genomic prediction for maize flowering time reveals novel insights into trait genetic architecture and improves prediction for breeding applications. bioRxiv doi: 10.1101/2025.07.15.664852.

Turner, B. M., 2009 Epigenetic responses to environmental change and their evolutionary implications. Philos Trans R Soc Lond B Biol Sci 364: 3403–3418.

van Eeuwijk, F. A., D. V. Bustos-Korts, and M. Malosetti, 2016 What should students in plant breeding know about the statistical aspects of Genotype × Environment interactions? Crop Sci 56: 2119–2140.

van Ittersum, M. K., P. A. Leffelaar, H. van Keulen, M. J. Kropff, L. Bastiaans, et al., 2003 On approaches and applications of the Wageningen crop models. Eur J Agron 18: 201–234.

Vasseur, F., L. Fouqueau, D. d. Vienne, T. Nidelet, C. Violle, et al., 2019 Nonlinear phenotypic variation uncovers the emergence of heterosis in Arabidopsis thaliana. PLOS Biol 17: e3000214.

Voss-Fels, K. P., M. Cooper, and B. J. Hayes, 2019 Accelerating crop genetic gains with genomic selection. Theor Appl Genet 132: 669–686.

Wade, M. J., 2002 A gene’s eye view of epistasis, selection and speciation. J Evol Biol 15: 337–346.

Wade, M. J. and C. J. Goodnight, 1998 The theories of Fisher and Wright in the context of metapopulations: when nature does many small experiments. Evolution 52: 1537–1553.

Werner, C. R., M. Zaman-Allah, T. Assefa, J. E. Cairns, and G. N. Atlin, 2025 Accelerating genetic gain through early-stage on-farm sparse testing. Trends in Plant Science 30: 17–20.

White, M. R., M. A. Mikel, N. d. Leon, and S. M. Kaeppler, 2020 Diversity and heterotic patterns in North American proprietary dent maize germplasm. Crop Sci 60: 100–114.

Wilkins, O., C. Hafemeister, A. Plessis, M.-M. Holloway-Phillips, G. M. Pham, et al., 2016 EGRINs (environmental gene regulatory influence networks) in rice that function in the response to water deficit, high temperature, and agricultural environments. Plant Cell 28: 2365–2384.

Wright, S., 1932 The roles of mutation, inbreeding, crossbreeding, and selection in evolution. In Proceedings of the Sixth International Congress of Genetics, pp. 356–366, Ithaca, NY, Brooklyn Botanic Garden.

Xing, K., H. Li, D. Kong, and C. Chen, 2023 Editorial: Plant responses to environmental stresses based on physiological and functional ecology. Front Plant Sci 14: 1290405.

Xiong, W., M. Reynolds, and Y. Xu, 2022 Climate change challenges plant breeding. Curr Opin Plant Biol 70: 102308.

Yabe, S., M. Yamasaki, K. Ebana, T. Hayashi, and H. Iwata, 2016 Island-model genomic selection for long-term genetic improvement of autogamous crops. PLOS ONE 11: e0153945.

Yu, X., X. Li, T. Guo, C. Zhu, Y. Wu, et al., 2016 Genomic prediction contributing to a promising global strategy to turbocharge gene banks. Nat Plant 2: 10.1038/NPLANTS.2016.150.

Zhao, Z., E. Wang, J. A. Kirkegaard, and G. J. Rebetzke, 2022 Novel wheat varieties facilitate deep sowing to beat the heat of changing climates. Nat Clim Chang 12: 291–296.

